# Data-Efficient Exploration of Enzyme Function Using Family-Specific Machine Learning

**DOI:** 10.64898/2026.06.02.729712

**Authors:** F. Hafna Ahmed, Asher Bender, Asiri Wijesinghe, Allen Zhu, Lu Zhang, Leigh Gebbie, Adrian Marsh, Chie Ishitate, William Holdsworth, Candice Jones, Andrew C. Warden, Helen Power, Cheng Soon Ong, Daniel Steinberg, Robert E. Speight

**Affiliations:** Environment, CSIRO, Clunies Ross Street, Canberra, ACT 2601, Australia; Technology, CSIRO, Clunies Ross Street, Canberra, ACT 2601, Australia; Advanced Engineering Biology Future Science Platform, CSIRO, Dutton Park, Brisbane, QLD 4102, Australia

## Abstract

Enzymes are essential biocatalysts across diverse industries, driving demand for high-performing variants. Foundation models are attractive for guiding enzyme discovery, but often lack the resolution to model subtle variations driving function within homologous families. Navigating these rugged functional landscapes to identify elite variants remains challenging and experimentally costly, even when guided by such models. Here we show that coupling dense, family-specific experimental screening with targeted, sequence-based deep learning provides a data-efficient discovery strategy. We experimentally screened 1,513 natural homologues from an esterase superfamily (>7,500 assays) and used this functional landscape to train task-specific models that predict activity, thermostability, and substrate specificity from sequence alone. Prospective experimental validation of previously untested sequences demonstrated that these task-specific models significantly outperformed generalist pre-trained and physics-based models in enriching for target traits. Residue-level attribution further indicated that the models captured sequence patterns consistent with underlying structural features. Finally, retrospective simulations showed that iterative retraining compresses the search space, discovering 60% of top-tier hits using nearly half the samples required by pre-trained baseline models. Together, these results highlight that machine learning can provide mechanistic insight, and that integrating targeted data acquisition with iterative machine learning provides a more data-efficient discovery strategy than relying on generic model scale.

## Introduction

Machine learning (ML) and artificial intelligence (AI) are increasingly shaping modern protein engineering workflows by enabling computational prioritisation of enzyme sequences for experimental testing. The rapid expansion of protein sequence databases has created unprecedented opportunities to explore functional diversity, yet only a small fraction of these sequences has been experimentally characterised^1^. As a result, there is growing interest in models capable of moving “from sequence to function” by predicting activity, substrate specificity or stability directly from primary sequence to reduce the burden of experimental screening^1–4^. Because enzyme fitness landscapes are often rugged, small sequence changes can have disproportionate effects on activity or stability ^5^, making high-performing variants difficult to identify through unguided search alone. Ultimately, the success of protein engineering hinges on identifying functional enzymes within realistic constraints, making effective sequence prioritisation a central challenge in enzyme discovery^4,6–8^.

Recent advances in large pretrained protein sequence models demonstrate that neural networks trained on expansive sequence datasets can extract statistical representations directly from amino-acid sequence and support tasks such as gene annotation^9^, enzyme property prediction and functional inference^10–12^, and structural prediction^13^. These approaches expand the scope of sequence-based modelling, enabling exploration of regions of sequence space where established similarity-based inference methods provide limited guidance^9,10^. However, general predictive performance does not automatically translate into task-specific discovery success. Models trained on highly heterogeneous datasets may generalise poorly to specific enzyme families and often fail to capture practical experimental constraints such as expression variability, assay context or condition-specific effects. Consequently, while pretrained representations on broad heterogeneous datasets provide powerful general-purpose features, their utility for targeted enzyme discovery remains uncertain.

A recurring challenge in ML-guided enzyme engineering lies in the nature, quantity and quality of available training data. Furthermore, existing labels are often heterogeneous, noisy or derived from indirect proxies^1,3^, and differences in assay design further introduce variability and complicate cross-dataset comparisons. As a result, reported model performance is highly sensitive to dataset construction and evaluation strategy, as demonstrated by systematic benchmarking studies^14–16^. Naïve train-test splits may inflate apparent accuracy by allowing closely related homologues to appear in both sets^14,15^, whereas extrapolation-focused or family-aware benchmarks reveal substantially reduced generalisation^15,16^. In practice, however, many enzyme discovery workflows prioritise functional variation within homologous families rather than extrapolation across distant sequence space. These observations help explain why models that perform well on curated retrospective benchmarks may struggle to enrich for improved variants in prospective discovery campaigns^2,6^.

Beyond data quality, these observations have prompted increasing focus on aligning modelling strategies with the experimental and evolutionary scope of enzyme discovery workflows. Large protein language models capture broad evolutionary information across protein space, yet their generality can limit sensitivity to subtle biochemical variation within a specific enzyme family^10,11^. In contrast, family-specific modelling strategies, including representation learning approaches such as embeddings trained on reconstructed ancestral sequences^17^, and supervised models trained within defined sequence landscapes ^18,19^, have been proposed to better capture local evolutionary structure within narrow sequence landscapes. It is evident that increasing model parameter counts or architectural complexity alone does not necessarily improve discovery outcomes, whereas effectiveness often depends on matching model scope with the diversity and scale of available experimental data^2,7,16^. Iterative or adaptive learning strategies, such as active-learning assisted directed evolution, further emphasise that predictive models become most effective when tightly integrated with experimental feedback and feasible screening workflows^2,7,20^. Furthermore, task-specific end-to-end predictive models allow for straight-forward application of interpretability and attribution methods, enabling deeper human understanding and verification of the discovered predictive relations^21^.

In practice, many enzyme discovery workflows still rely on similarity-based frameworks to navigate protein families when experimental data are unavailable, including automated annotation pipelines implemented across major protein resources^22–24^. Conventional sequence-based enzymology approaches such as phylogenetic reconstruction, motif-based annotation, and sequence similarity networks, provide effective frameworks for organising evolutionary relationships and guiding representative sampling ^25,26^. These methods describe how sequences relate to one another but offer limited insight into comparative biochemical performance across homologues. Together, these observations highlight a broader challenge in enzyme discovery, where mapping evolutionary relationships across sequence space does not necessarily enable reliable prioritisation of targets within functionally diverse families.

Esterases provide a compelling system in which to examine these challenges. They are widely used across industrial biotechnology due to their catalytic versatility, broad substrate scope and adaptability to immobilised formats ^27^. Within this class, the SGNH hydrolase superfamily remains underexplored in comparison to classical α/β-hydrolases. SGNH esterases share a conserved Ser-Gly-Asn-His catalytic tetrad yet exhibit substantial diversity in substrate preference and biochemical behaviour^28,29^. PpEST is an esterase within this superfamily that hydrolyses polyester substrates^30^. We recently explored a sub-family of PpEST-related proteins, consisting of sequence-divergent enzymes that share polyesterase function but vary widely in catalytic efficiency, stability, and expression properties^31^. Although extant protein characterisation and structure-guided engineering have developed the catalytic potential of PpEST family proteins^31,32^, homology alone provides limited guidance on process-relevant traits. This combination of conserved function and heterogeneous biochemical performance makes the PpEST family of SGNH esterases a useful model system for evaluating how sequence-based ML performs under realistic discovery conditions.

In this study, we combined large-scale experimental characterisation with sequence-based deep learning to map functional diversity within the PpEST family of SGNH esterases (**Figure 1**). We experimentally assayed 1,513 wild-type homologues across three substrates and measured thermostability at two temperatures, generating a detailed functional landscape within a single enzyme family. Using this dataset, we trained task-specific convolutional neural networks (CNNs) to predict activity, thermostability, and substrate preference. We then directly compared their performance to general pretrained models that lacked family-level retraining. Family-specific models more accurately captured observed biochemical variation and enabled prospective identification of improved variants among previously untested homologues. Feature attribution analysis further revealed structurally coherent regions associated with functional divergence, while retrospective simulations demonstrated that iterative retraining improves discovery efficiency under limited screening budgets. Together, these results show that experimentally grounded, family-level models can outperform general pretrained models and provide a data-efficient strategy for prioritising functional diversity within homologous enzyme families.

**Figure 1.**
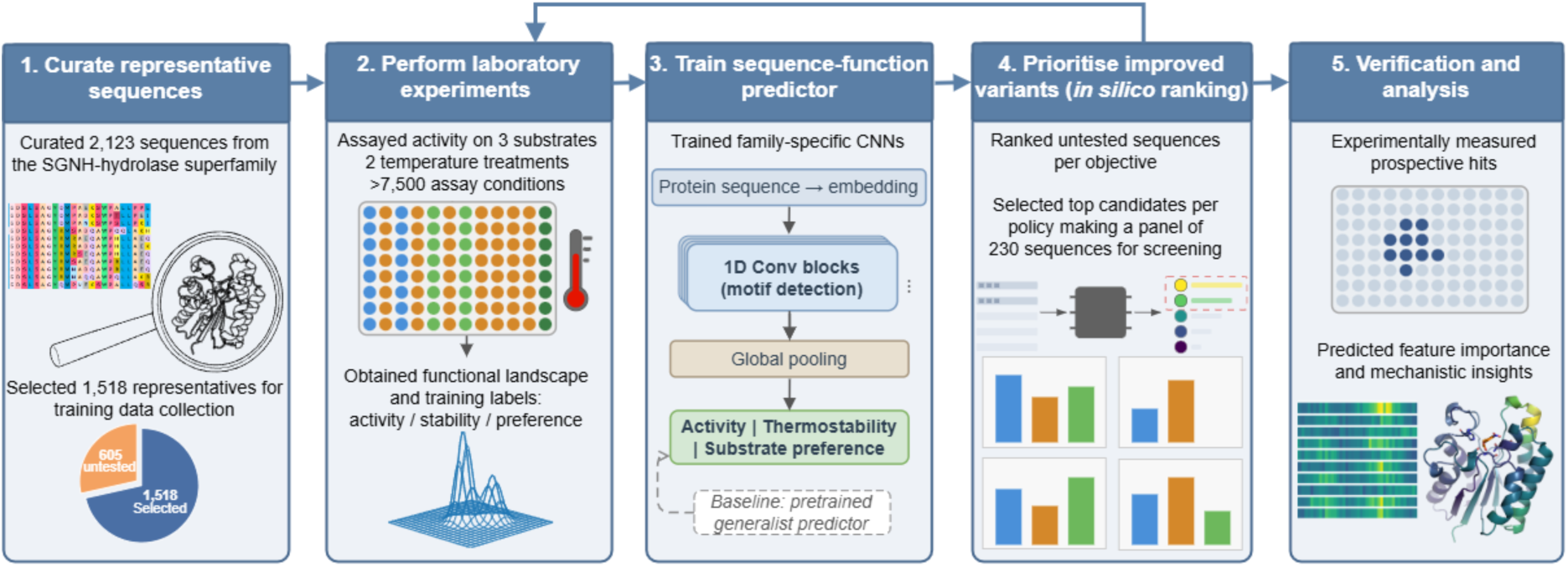
Machine learning–guided enzyme discovery workflow. Top (dark blue panels) shows our recommended workflow for integrating biological screening with sequence-based machine learning. Representative sequences are curated from a homologous family, experimentally assayed, used to train sequence–function predictors, ranked *in silico*, and selected variants experimentally verified. If required, highest ranked sequences are selected for another round of assays, providing better training data, leading to more accurate predictions. Feature attribution identifies sequence regions associated with model predictions. The lower panels show the implementation in this study. A total of 2,123 SGNH-hydrolase homologues were curated, of which 1,518 enzymes were initially experimentally assayed towards three substrates and two temperature pre-treatments (>7,500 assays). Convolutional neural networks were trained to predict activity, thermostability and substrate preference from sequence alone. The models ranked the remaining 605 untested sequences for different attributes (activity, thermostability and substrate preference), from which 230 candidates were selected for experimental validation and feature importance analysis.

## Results

### Mapping sequence and functional diversity of PpEST family esterases

We previously identified and curated the PpEST-like polyesterase family, comprising 2,123 homologous from the broader SGNH-hydrolase superfamily^31^. From our previous sequence similarity network (SSN) analysis, 1,987 of these sequences cluster tightly with PpEST, while the remainder form a closely related sister clade. From the 1,987 PpEST-like sequences, we selected a non-redundant subset of 1,518 representatives for initial experimental characterisation (pair-wise sequence identity 20-90 %, median 40 %, **Figure S1**). Automated signal-peptide truncation also introduced additional variation, with some sequences retaining extra N-terminal residues that may affect protein stability and function^31^. This curated subset is sufficiently large to capture natural functional diversity within a homologous family, enabling systematic comparison of biochemical variation across related enzymes while remaining experimentally tractable for high-throughput characterisation.

To interpret the organisation of this space, we generated *sequence-only* embeddings using the ESM-2 language model^33^ and used an unsupervised model to group sequences into 15 clusters. Visualising these representations via 2D projections revealed a highly structured landscape (**Figure 2A**), where the embedding-based clustering was coherent with taxonomic grouping (**Figure 2B**), and the uncharacterised sister clade formed a distinct separated cluster (cluster 3, **Figure 2A**). Colouring a phylogenetic tree with these clusters demonstrated a strong correspondence between embedding-based clustering and phylogenetic branching (**Figure 2C**). Even though the embeddings and clusters were generated from primary sequences without any explicit taxonomic, structural, or phylogenetic information, they broadly capture the major evolutionary lineages within the family.

**Figure 2:**
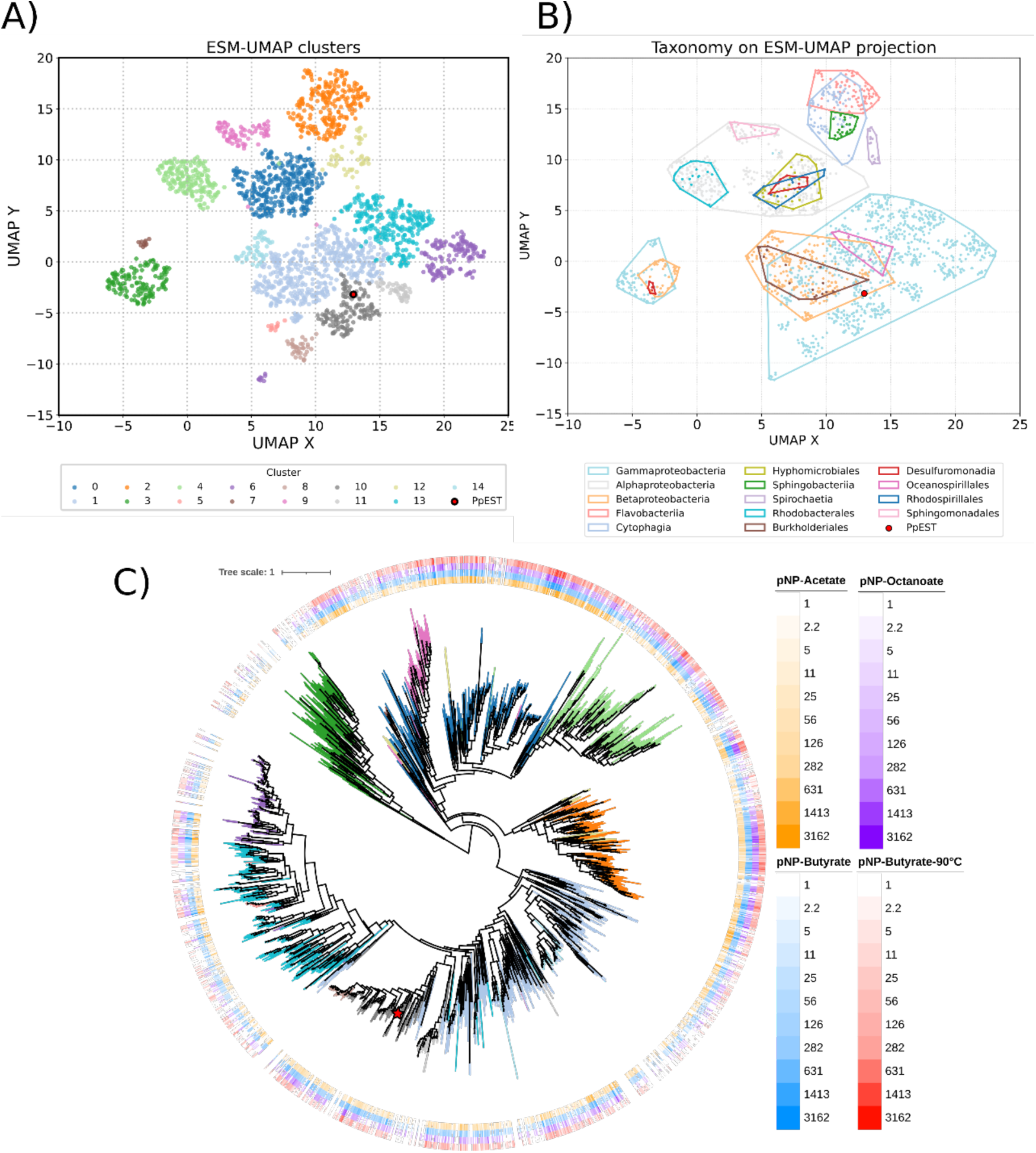
Sequence similarity and structural organisation of the PpEST-like polyesterase family. In **A** and **B**, each point represents a 2D UMAP projection of the ESM-2 embeddings for the sequences that were curated in our previous work ^31^. **A.** Points are coloured by the spectral clusters derived from a 50-dimensional UMAP manifold. Cluster 3 represents an experimentally uncharacterised clade excluded from the initial dataset. **B.** Points are coloured based on the taxonomy of the originating organism. **C.** Phylogenetic tree of the sequences rooted at cluster 3, with branches coloured based on the clustering identified in panel **A.** Log maximum velocities (Log MaxV (mOD/min) for the three substrates pNP-acetate, pNP-octanoate, and pNP-butyrate before and after heating at 90 °C are shown for all 1,743 sequences characterised in this study.

For experimental characterisation, the 1,518 representatives were expressed in *Escherichia coli* and screened in a high-throughput workflow for esterase activity, substrate preference and thermostability (**Figure 1, Table S1**). Activities were measured directly from clarified lysates, providing an integrated readout of catalytic performance and expression level. This format enabled reproducible comparison of large numbers of homologues, capturing relative variation in enzyme performance across the family rather than enzyme turnover for individual variants, while incorporating expression-related effects that influence practical enzyme utility.

To assess overall activity and substrate preference, we used three *p*-nitrophenyl (pNP) esters: pNP-acetate (C₂), pNP-butyrate (C₄), and pNP-octanoate (C₈). These substrates are soluble under aqueous, high-throughput conditions and report esterase activity *via p*-nitrophenol release monitored spectrophotometrically^31^. pNP-butyrate is the optimal substrate for PpEST homologues^30,31^, while pNP-octanoate and pNP-acetate were used to capture variability in chain-length specificity. Substrate and lysate concentrations were validated within the linear detection range (**Figure S2**), and assay reproducibility was confirmed using controls and 40 enzymes repeatedly measured across independent plates (**Figure S3, Figure S4**). The final dataset comprised measurements for 1,513 sequences, excluding one homologue that failed transformation and four that did not pass quality control in all assays.

High-throughput screening against the three substrates (n = 3 on average) yielded broad and reproducible activity distributions (**Figure 2C, Figure S5**), with maximum velocity (MaxV) values spanning over three orders of magnitude. As previously observed^30,31^, pNP-butyrate generally produced the highest mean activity for each enzyme, followed by pNP-acetate then pNP-octanoate (**Figure 2C, Figure S5-S7**). Most enzymes exhibited low to moderate activity (<500 mOD/min) and standard deviations varied across the dataset (**Figure S5**), with activity also varying across UMAP and taxonomic clusters (**Figure 2C, Figure S6-S7**).

Pairwise comparisons of MaxV across substrates revealed clear trends in chain-length preference (**Figure S8-S9**). Correlation between pNP-acetate and pNP-octanoate was weakest (ρ=0.66), suggesting that enzymes tend to specialise for either short- or long-chain esters but rarely both. In contrast, pNP-acetate and pNP-butyrate activities were more strongly correlated (ρ=0.81). MaxV rank comparisons reinforced this trend, with pNP-acetate and pNP-butyrate preferences correlating more closely than either substrate with pNP-octanoate (**Figure S10**). This pattern is consistent with the closer chemical similarity between pNP-acetate and pNP-butyrate, which differ by only two carbons, whereas the longer acyl chain of pNP-octanoate likely imposes distinct constraints on substrate accommodation.

Thermostability was evaluated by measuring residual pNP-butyrate activity after lysates were incubated at either 60 °C or 90 °C for 10 minutes, which is an effective initial screen for identifying thermostable enzymes^31,34^. The percentage retention values were strongly correlated between the two temperatures (ρ=0.88), with several enzymes showing both high overall activity and high residual activity after heat treatment (**Figure S8, Figure S10**). Because activity was measured in clarified lysates, protein expression contributes to the magnitude of MaxV and thus to the measured thermostability. Comparison of MaxV values with SDS-PAGE band intensity for 39 enzymes revealed a generally positive but dispersed relationship (**Figure S11**), with activities varying among enzymes with similar expression levels. This indicates that while expression level contributed to measured activity, it did not fully explain the observed variation in activity or thermostability. Thermostability also showed no clear correlation with pNP-octanoate preference relative to pNP-butyrate (**Figure S12**), indicating that apparent thermostability and substrate preference are largely independent traits within this enzyme family.

### Sequence-only ML predictions of activity, thermostability and substrate specificity

To determine whether the biochemical variation observed across the 1,513 SGNH-hydrolase homologues could be inferred directly from amino-acid sequence, we developed supervised machine-learning models trained exclusively on the curated assay data (**Figure S13**). These models were designed to predict three target traits: overall activity, thermostability, and octanoate preference (see Methods).

Convolutional neural network (CNN) predictors were trained on the initial screen data. As is typical for complex biological regression problems, the hold-out validation coefficients of determination (R²) were modest for these predictors across the tasks (**Table 1**). However, the models successfully captured rank-order performance. Predictions were strongest for thermostability (Spearman ρ ≈ 0.64–0.66) and octanoate preference (ρ ≈ 0.60), whereas overall activity showed more moderate predictability (ρ ≈ 0.47).

**Table 1.**
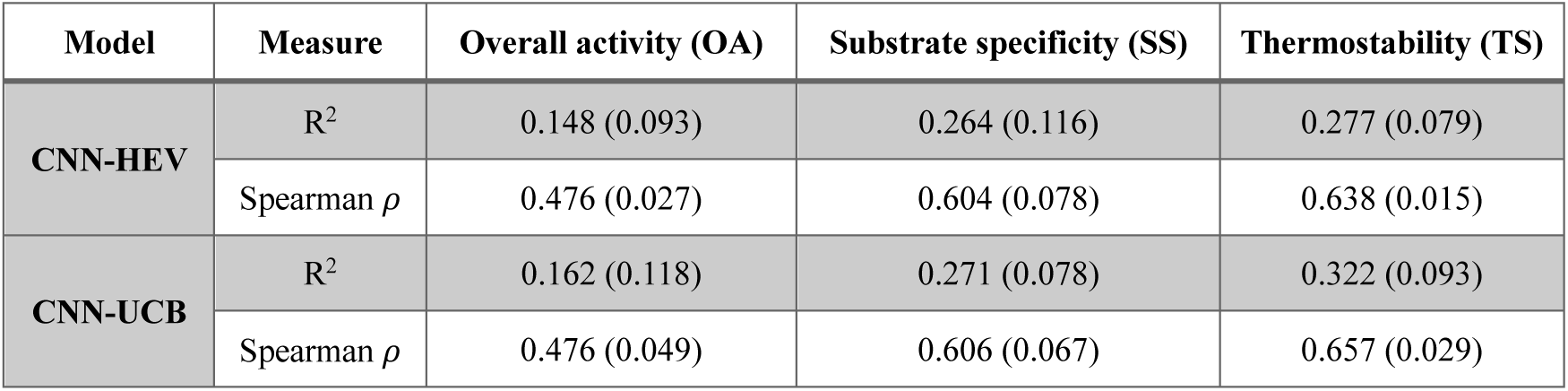
Out-of-fold prediction performance for the CNN-based models on the different prediction tasks. We report ±1 std. dev. in brackets from the five folds.

| Model | Measure | Overall activity (OA) | Substrate specificity (SS) | Thermostability (TS) |
| --- | --- | --- | --- | --- |
| CNN-HEV | $R^2$ | 0.148 (0.093) | 0.264 (0.116) | 0.277 (0.079) |
| | Spearman $\rho$ | 0.476 (0.027) | 0.604 (0.078) | 0.638 (0.015) |
| CNN-UCB | $R^2$ | 0.162 (0.118) | 0.271 (0.078) | 0.322 (0.093) |
| | Spearman $\rho$ | 0.476 (0.049) | 0.606 (0.067) | 0.657 (0.029) |

Two selection strategies were then used to prioritise candidates from the remaining 605 untested sequences. The first strategy selected sequences based on their highest expected value from a single CNN (CNN-HEV). The second strategy was based on an upper confidence bound from an *ensemble* of randomised CNN predictors (CNN-UCB). The CNN-HEV strategy is an entirely greedy (or exploitative) method whereas CNN-UCB is designed to balance exploration and exploitation^35^.

In addition to the supervised sequence-only models, we evaluated published computational methods as baseline selection procedures for all three experimental tasks (**Table S2**). These consisted of structure-based molecular docking and eight zero-shot machine-learning models trained on broad, multi-family datasets (see Methods). Because the models differed in their trained tasks, we selected predictors that positively correlated with experimental results and combined them via rank aggregation to obtain a single ranked candidate list per target.

To prospectively validate these strategies, we generated a deduplicated experimental panel of 230 untested sequences. To ensure a fair comparison, we allocated equivalent selection budgets across the methods, pooling up to 50 top-ranked candidates per task from each strategy (CNN-HEV, CNN-UCB, and pre-trained baselines), alongside 30 stratified random sequences to provide a baseline control across the sequence space. The two CNN strategies showed strong agreement, nominating 32 common candidates for overall activity and 21 for both thermostability and octanoate preference (**Figure S14**). In contrast, the pretrained models showed limited overlap with the CNN candidates, indicating that family-specific and generalist models prioritise fundamentally different sequences.

Projection of the selected candidates onto the 2D sequence embedding projection revealed task- and policy-dependent selection patterns across the protein landscape (**Figure 3**). For all three tasks, CNN-HEV and CNN-UCB selections were broadly concentrated in regions enriched for high-performing training examples, showing similar spatial selection patterns. The UCB strategy further expanded coverage (exploration) relative to HEV, incorporating candidates from less densely represented clusters. In contrast, random and pretrained selections were more diffusely distributed and showed less clear enrichment within regions associated with strong experimental performance. The pre-trained models also more frequently prioritised sequences within the related sister clade (with no training data) when predicting overall activity and octanoate specificity, whereas CNN-HEV and CNN-UCB generated confident predictions in this region primarily for thermostability.

**Figure 3.**
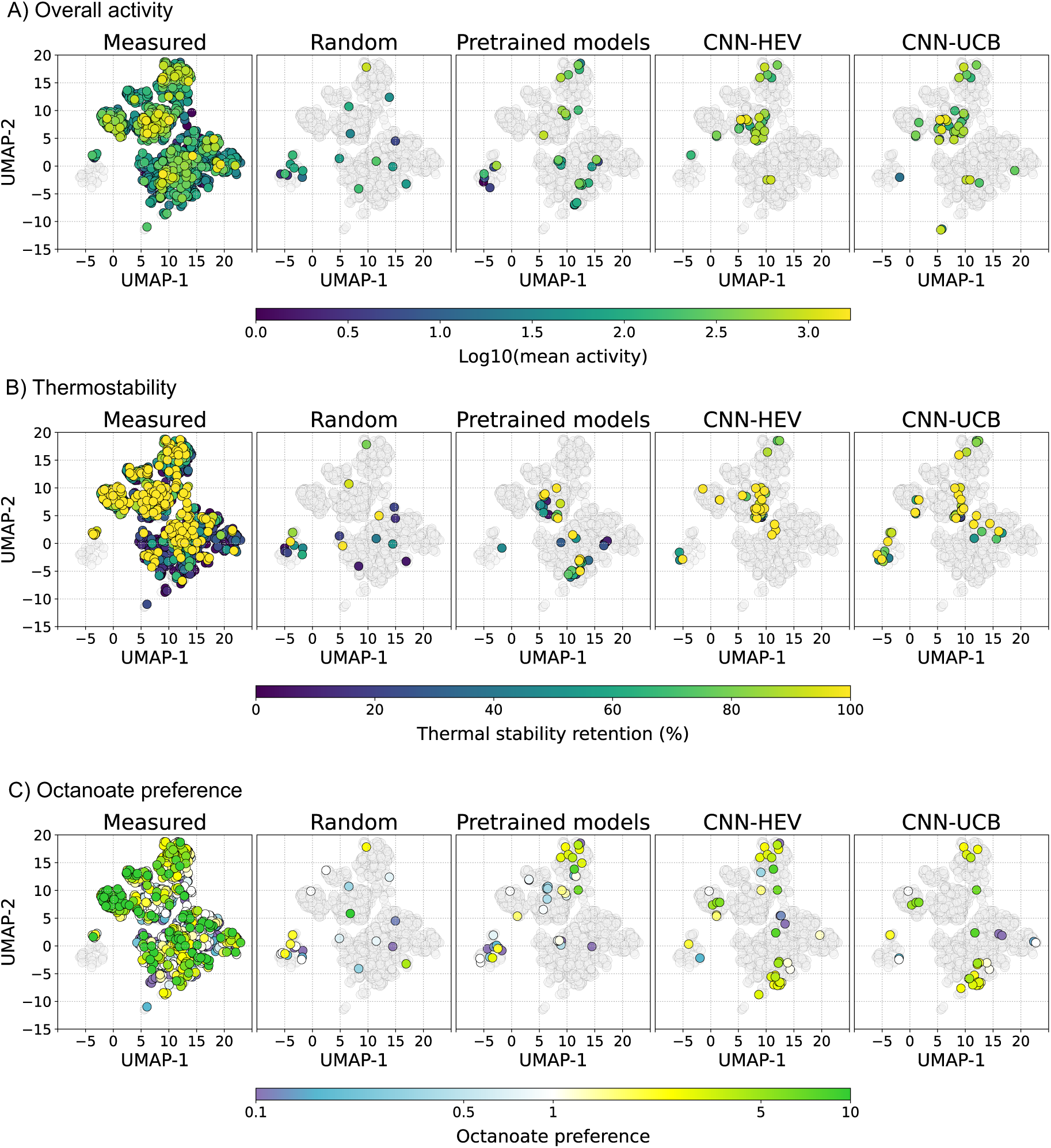
Sequence-function landscape of initial and prospective cohorts. Two-dimensional projections of the sequence embeddings (as in Figure 2) coloured by experimentally measured outcomes from the initial screening cohort (leftmost column) and the prospective validation cohort (remaining columns). The initial cohort comprises the 1,513 sequences used for training (uncharacterised sequences shown in grey); the validation cohort comprises sequences selected from the uncharacterised pool using random selection, pretrained models, CNN-HEV, and CNN-UCB policies (entire PpEST-like polyesterase family shown in grey). Rows represent: **(A)** overall activity, **(B)** thermostability, and (C) octanoate preference. For visual clarity, thermostability and octanoate preference ratios are clamped to 0–100% and 0.1–10, respectively.

### Prospective experimental validation of ML-guided sequence selection

To determine whether these distinct *in silico* selection patterns translated to functional improvements *in vitro*, we synthesised, expressed, and screened the 230 candidate sequences under identical assay conditions (**Figure 3**, **Figure S15**). For all tasks, both CNN strategies demonstrated highly significant positive distributional shifts (p < 0.001) compared to the original screening baseline (**Table S3**). In contrast, the pre-trained models (p ≥ 0.08) failed to provide significant positive distributional shifts, reflecting their broad and family-agnostic training objectives. Both CNN strategies also consistently achieved higher hit rates for top-tier variants than the pre-trained models across all prediction targets (**Table S4**). This enrichment was most pronounced and statistically significant for the overall activity task, yielding a multi-fold increase in hits. While the CNN strategies reliably improved the bulk population across all tasks, they did not significantly enrich for thermostability and octanoate preference. This indicates that capturing broad functional gradients is an easier task than locating the rugged fitness peaks that define rare elite variants.

Mapping the experimental outcomes onto the sequence landscape represented by the 2D sequence-embedding projections revealed that high-performing CNN selections localised within, or adjacent to, regions already enriched for strong performers in the training data (**Figure 3**). Furthermore, CNN-UCB showed a broader spatial diversity of high-performing selections across the landscape compared to CNN-HEV. This increased diversity is consistent with the UCB strategy, which uses disagreement among ensemble members to balance exploitation with exploration of sequences with high predicted performance and variance. Importantly, several highly thermostable enzymes originated from the uncharacterised sister-clade to PpEST lacking any prior representatives, demonstrating the CNN models’ ability to discover functional variants beyond conventional similarity-based screening. In contrast, sister-clade variants were rarely recovered among selections for overall activity or octanoate preference, suggesting that sequence features associated with thermostability are more transferable across related enzyme families.

Overall, these results suggest that supervised sequence-only models substantially increase the discovery yield of functional enzymes from within large unlabelled sequence pools, with ensemble and uncertainty-driven selection outperforming random and pretrained model-based strategies.

### Structural interpretation of residue predictive importance

To examine which sequence features most strongly influenced model predictions, we performed per-residue predictive importance analysis using the CNN ensemble across the three prediction tasks: overall activity, thermostability, and octanoate preference. We focused on cluster 2 as a case study (**Figure 2A**), which contains 214 sequences experimentally characterised in the initial screen that generally showed strong performance across the three tasks (**Figure S6**). Additionally, the CNN ensemble predicted several sequences within this cluster to be high performing (**Figure 3**).

The residue predictive-importance heatmaps on the multiple sequence alignment (MSA) of cluster 2 showed clear task-specific patterns (**Figure 4A, Figure S16**). Across sequences, regions of positive or negative importance formed clear bands rather than isolated residue-level signals, indicating that the model relied on reproducible and structured sequence patterns. These patterns were consistent across sequences for each task, albeit skewing towards more negative importance values in the bottom 10 sequences compared to the top 10 sequences when ranked by predicted MaxV (**Figure 4A**), as expected for the predictive importance technique used^21^. The regions of highest residue importance in terms of either positive or negative contribution are also differentially conserved in amino acid composition for the different tasks (**Figure 4C**), suggesting that increased or decreased amino acid variation is probably not the basis for the importance scores.

**Figure 4.**
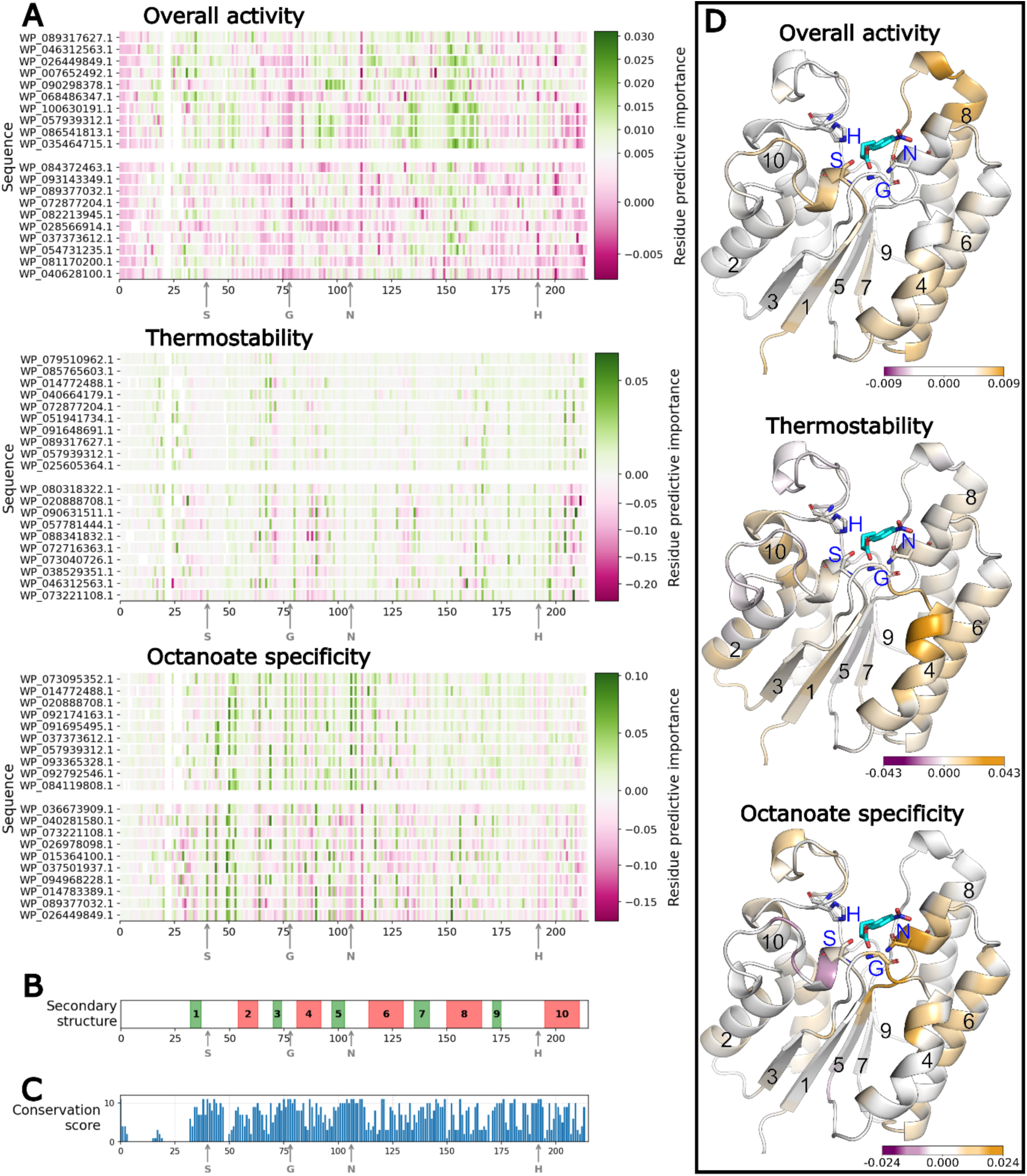
Prediction of residue importance in the protein sequences used for prediction by the CNN-UCB ensemble model. **A.** Multiple sequence alignments (MSA) of the top and bottom 10 sequences from each of the three tasks (overall activity, thermostability, octanoate preference) from cluster 2 in Figure 2A, ranked by predicted MaxV. Individual amino acids are coloured by their residue predictive importance that reflects contribution magnitude to the prediction, where sum importance is approximately equivalent to predicted MaxV (**Figure S16**). Sequence IDs are shown on the Y-axis and the positions of the catalytic S, G, N, H residues in the alignment are shown on the X-axis. **B.** Secondary structure prediction for the MSA, where beta-sheets are in green and helices are in pink. **C.** The conservation score for the MSA calculated by Jalview^36^, where higher values indicate higher conservation. **D.** The normalised average feature importance for sequences in cluster 2 (122 for overall activity, 100 for thermostability, 108 for octanoate preference), mapped onto the Alphafold3^37^ structure prediction for WP_015364100.1 that had no gaps in the MSA. The colour gradient reflects per-residue importance values normalised within each sequence to sum to 1 and then averaged across the MSA at each aligned residue position. This highlights sequence positions that consistently contribute to model predictions across the cluster, independent of differences in overall activity. Secondary structure features are labelled to match panel B, and the S, G, N, H catalytic residues are labelled and shown as sticks. PNP-butyrate is shown as a substrate in cyan.

Although predictive importance does not imply true causal mechanism, the contrasting heatmap patterns indicate that the biological traits evaluated in the three different tasks are linked to different sequence regions of the enzyme (**Figure 4A**, **Figure 4B**). To visualise the spatial distribution of these regions, we mapped the average normalised residue predictive importance across the cluster for each task onto the Alphafold3 structure of the same representative sequence (**Figure 4D**). For overall activity the highest attributed regions include α-helix 8 and the preceding loop region that forms one side of the distal portion of the substrate-binding groove, along with the catalytic serine. These regions could modulate substrate access and positioning, which can influence catalytic performance. In contrast, octanoate specificity highlighted the core catalytic site, including the loop adjoining the catalytic serine, the catalytic asparagine and adjoining loop, as well as the catalytic glycine and a portion of the adjacent α-helix 4. These elements are positioned to influence the geometry of the active-site region and can determine local active-site architecture. For thermostability, rather than the catalytic centre or substrate binding groove, the highlighted residues are mostly distributed among the surface α-helices 4, 2, 6, and 10, that are likely to contribute to the structural scaffold and thus stability of the enzyme.

Taken together, these analyses show that the sequence-only CNN ensemble model captures distinct, task-specific patterns that map onto interpretable structural features, highlighting regions that can be associated with predicted biochemical outcomes. This mapping offers a complementary perspective for interpreting sequence–structure–function relationships within this enzyme family.

### Iterative retraining compresses the experimental search space

To evaluate how different sequence selection strategies convert limited screening budgets into functional discoveries, we simulated an offline campaign using the combined dataset (1,743 sequences) as a ground-truth functional landscape. We compared adaptive strategies that iteratively retrained on new observations (CNN-HEV and CNN-UCB) against static baselines (random sampling and pre-trained models). Performance was assessed using cumulative hit rate, recall, and cumulative regret, with a “hit” defined as an enzyme ranking in the top 10% for a given task.

Across all three tasks (overall activity, thermostability, and octanoate preference) the adaptive CNN strategies demonstrated a clear early advantage over both the random baseline and pretrained models (**Figure 5**). Their cumulative hit rate (**Figure 5**, top row) peaked within the first 300 samples, achieving a hit density nearly double that of the random baseline. This early peak illustrates an efficient selection effect, where the models successfully identify the most promising candidates first. As the pool of available high-performers is depleted, the hit rate naturally converges toward the baseline. The pretrained models only offered marginal improvements over random sampling, suggesting that broad, zero-shot biological knowledge is a poor substitute for the family-specific active learning.

**Figure 5.**
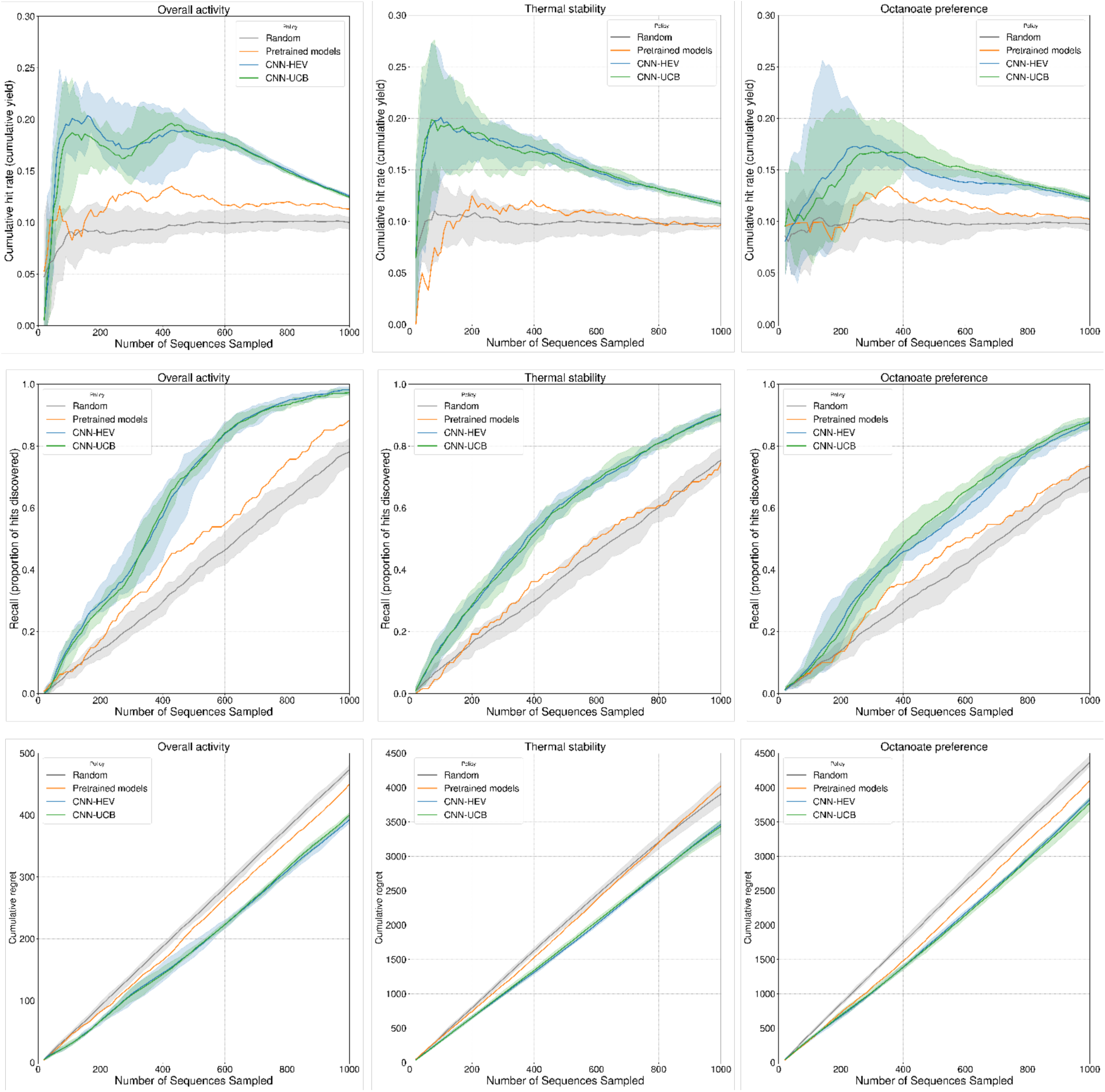
Simulated iterative sampling of the PpEST functional landscape. Columns represent the three screening tasks: overall activity (left), thermostability (middle), and octanoate preference (right). Rows display different performance metrics: cumulative hit rate (top), proportion of hits discovered (middle), and cumulative regret (bottom). Four selection strategies are compared: random selection (grey), pretrained models (orange), CNN-HEV (blue), and CNN-UCB ensemble (green). Simulations were performed in batched rounds, with each strategy selecting a batch of unobserved sequences per iteration. Each strategy began with identical starting data and was evaluated over independent runs with different random seeds. Shaded regions represent the distribution of results across these runs; the pretrained model’s strategy (orange) is shown only as a single line as it follows a deterministic static ranking.

The recall curves (**Figure 5**, middle row) provide the most direct evidence of search space compression. In the overall activity task, both CNN strategies discovered 60% of all available hits after approximately 400 samples. To achieve an equivalent discovery yield, the random sampling strategy requires over 750 sequences, representing over 1.875 times the experimental effort. In the most challenging task, octanoate preference, both CNN strategies successfully captured at least 45% of all available hits within the first 400 sequences sampled. While the discovery yield is lower than the simpler, overall activity task, random sampling still requires over 650 sequences to reach the same yield - representing over 1.625 times the experimental effort.

The cumulative regret curves (**Figure 5**, bottom row) quantify the total functional performance foregone through sub-optimal selections. Both CNN strategies accumulated regret at a lower rate than the baselines. This performance gap indicates that the adaptive models were better at prioritising high-performing variants throughout their sampling trajectory. In a physical campaign, this sustained ability to steer laboratory resources away from poor-performing sequences, translates into minimised experimental waste.

While the specific discovery rates in this simulation depend on the functional landscape of the PpEST family, the results support a broader advantage for adaptive, model-in-the-loop exploration of family-wide sequence spaces. By integrating iterative experimental feedback into the selection process, such strategies can increase discovery efficiency while reducing wasted experimental effort.

## Discussion

Protein engineering is inherently a multi-objective optimisation problem. To be industrially or therapeutically useful, an enzyme must often satisfy multiple biochemical constraints simultaneously, including maintaining high catalytic turnover while exhibiting thermostability and precise substrate specificity. Functional proteins exist as rare, isolated islands within a rugged fitness landscape, where even minor mutational perturbations can dismantle the epistatic networks required for catalysis or stability. Consequently, the probability of identifying elite variants through random sampling or naive library generation remains low^5,6^. Recently, large protein language models have expanded our ability to navigate sequence space, enabling advances in structure prediction and sequence generation^12,33,37^. However, translating these capabilities into reliable discovery of high-performing variants within defined enzyme families remain challenging.^6,16^

To bridge this gap, we must reconsider how we sample sequence space. Existing data acquisition strategies typically occupy two extremes: deep mutational scanning and large-scale database mining. Deep mutational scanning densely samples variants around a single scaffold but remains constrained to local sequence neighbourhoods, limiting its ability to capture broader evolutionary diversity. Conversely, large-scale database mining samples broadly across global protein space but often involves proteins that are too evolutionarily distant to resolve the determinants of biochemical performance within a specific enzyme family. Neither approach adequately captures the intermediate regime of family-specific diversity where homologous enzymes diverge functionally while retaining sequence and structural similarity. Recent family-level representation learning approaches similarly emphasise the value of modelling within constrained evolutionary landscapes, although these have primarily focused on representation rather than prospective validation^17^. Our study occupies this critical “middle ground”, where we curated a detailed dataset that documents the functional landscape of a single homologous family. Importantly, our dataset contains low-, average-, and high-activity variants, populating the training data to enable models to learn features that distinguish poor, average or elite performers, and to resolve fine-grained structure-function relationships. This scale of diversity differs from many enzyme discovery datasets and enabled broad sampling of within-family functional diversity while remaining compatible with experimental characterisation. Our detailed data enabled us to test a “Functional Fidelity” hypothesis: that for specific biophysical predictions, the consistency, relevance and density of training data outweigh the scale of the model architecture.

Our results strongly support this hypothesis. Simple, task-specific CNNs trained on our data consistently outperformed zero-shot predictions from pre-trained models, consistent with recent observations that model performance depends strongly on alignment between training data and application domain^7,16^. Hence, for defined engineering campaigns, predictive success seems to depend less on the sheer size of the model architecture and more by the density and relevance of the experimental landscape used for training.

A recurring misconception in protein engineering is the expectation that predictive models can serve as oracles capable of near-perfect accuracy. Our findings suggest a more pragmatic philosophy: models are best utilised as imperfect filters designed for bulk enrichment rather than individual precision. The goal is not to replace experimental screening, but to compress the search space upstream of validation and improve efficiency. Accordingly, while our models did not yield a 100% success rate, the CNN-selected panels exhibited a “hit rate” nearly double that of random selection or zero-shot baselines. By enriching the sampled pool for high-performing variants, the models ensured that limited experimental capacity was concentrated on the most promising regions of sequence space. This demonstrates that even simple models, when trained on relevant local data, can serve as efficient enrichment filters that distinguish the peaks of high performance from the background of non-functional or average variation.

Our retrospective simulations reveal that the efficiency of this filtering process is maximised through iteration, with adaptive strategies substantially outperforming static one-shot predictions, recovering 60% of top-tier variants with 1.8-fold fewer samples. Rather than relying on a fixed initial ranking, an adaptive approach allows the model to correct its search trajectory in real-time, focusing experimental effort on increasingly enriched pools of candidates. This supports that a highly productive workflow for enzyme discovery is iterative: a cycle of prediction, measurement, and retraining that minimises “experimental regret” and consistently steers laboratory resources toward the most informative regions of the fitness landscape.

A central question arising from this work is whether sequence-based machine-learning models serve solely as local interpolators, or whether they can capture transferable biophysical rules that permit extrapolation. Our prospective validation supports the latter: we successfully identified functional, thermostable variants within the uncharacterised “sister clade”, a related lineage absent from our training data. Although these sequences are evolutionarily distinct, they remain within the broader SGNH-hydrolase fold, suggesting that family-specific learning can capture sequence determinants of stability and activity that generalise across evolutionary sub-lineages. This highlights the utility of dense, local landscapes not just for refining known variants, but for exploring the functional boundaries of a homologous group.

Beyond prediction, our feature attribution analysis challenges the persistent notion of deep learning as an impenetrable “black box.” We observed spatially coherent attribution patterns where distinct sequence regions correlated with specific functional traits. For instance, features associated with overall activity mapped to distal substrate-binding surfaces, while octanoate preference was linked to the local active-site architecture. Crucially, these attribution maps aligned with structural features despite the models being trained exclusively on primary (unaligned) sequence. This indicates that family-specific sequence models can implicitly learn patterns that encode functionally relevant constraints that reflect the underlying 3D structure even without explicit structural modelling or docking.

In this context, machine learning serves not only as a predictive engine but as a tool for knowledge discovery. By highlighting regions where variation drives performance, the models provide a data-driven framework for hypothesis generation, transforming it from a mere selection tool into a potential vehicle of biological insight. This reinforces a “human-in-the-loop” philosophy, where machine-learning does not replace mechanistic understanding but complements it by revealing sequence-structure relationships that may not be apparent from structural inspection alone.

While our results highlight the benefits of an integrated strategy of iterative, family-specific data generation and modelling, this approach is not without limitations. The “bespoke” nature of our approach represents a trade-off between accuracy and universality. Unlike foundation models that can be applied zero-shot to any protein sequence, our CNNs are scoped strictly to the SGNH-hydrolase family. Applying this framework to a different enzyme class would require the generation of a new, high-quality “seed” dataset. Given the demonstrated efficiency gains, we argue that this initial experimental investment is justified when targeting specific enzyme families or complex functional traits that remain elusive to generalist models.

Our study demonstrates that data-efficient enzyme discovery is driven less by the scale of the model architecture and more by the consistency, relevance and density of available training data. By coupling representative, high-quality data with active, local learning methods, screening campaigns can adapt to new information and refine their search trajectories in real time. This framework offers a scalable blueprint for enzyme discovery, demonstrating that intelligent, iterative data acquisition can unlock functional diversity more effectively than naive screening efforts or generalist models.

## Methods

### Generation of the experimental benchmark dataset

#### Sequence selection and analysis

In our previous work^31^, protein sequences homologous to the polyesterase PpEST from *Pseudomonas oleovorans* were retrieved from the NCBI RefSeq database and clustered using sequence similarity networks (SSNs) to define the PpEST-like clade (2,123 sequences) within the SGNH-hydrolase superfamily. From these sequences, 1,987 sequences cluster with PpEST, while the remaining form a closely related, but functionally uncharacterised cluster. From these, 1,518 non-redundant sequences were selected using CD-HIT^38^ at 89.5% sequence identity. Signal peptides were predicted using SignalP 6.0^39^ and truncated according to automated predictions without manual adjustment to ensure an unbiased and reproducible workflow.

Sequence embeddings were generated for all 2,123 sequences using the ESM-2 transformer model (650M parameters) ^33^. To obtain fixed-length numerical representations, the mean of the last four hidden layers was extracted, and mean pooling was applied across the sequence length (excluding padding and special tokens). These pooled representations were subsequently L1-normalised. Following L1-normalisation, the resulting 1,280-dimensional representations were reduced to a 50-dimensional manifold using UMAP^40^ (cosine distance metric, 15 nearest neighbours, minimum distance of 1). This dimensionality reduction mitigates the curse of dimensionality by minimising vector sparsity and filtering uninformative representation noise. The lower-dimensional manifold is a denser, more semantically meaningful neighbourhood graph for subsequent clustering. To group sequences based on their representational similarity, spectral clustering (using a nearest-neighbours affinity matrix) was then applied to this 50-dimensional manifold to partition the dataset into 15 clusters. Finally, to visualise the sequence landscape, the original 1,280 dimensional representations were projected down to a 2D representation using UMAP and overlaid with their corresponding cluster assignments and taxonomic metadata.

For phylogenetic analysis, all 2,123 sequences were aligned using MAFFT-DASH ^41^ and the tree was generated with IQ-TREE ^42^ using the LG+F+R9 evolutionary model. UltraFast Bootstrap (UF-Boot) support values and SH-aLRT branch test values were calculated for 1,000 replicates and the tree was visualised and annotated using the Interactive Tree of Life (iTOL) webtool^43^.

#### Plasmid and library design

DNA sequences were codon-optimised for *Escherichia coli* expression and synthesised with a C-terminal His₆-tag. Genes ordered from TWIST were cloned between the *NdeI* and *XhoI* sites of the T7-driven expression vector pET-29b+. The resulting plasmids were used to transform T7 Express lysY Competent *E. coli* (New England Biolabs) using a miniaturised heat-shock protocol (42 °C, 10 s, 1 µL cells + 100 nL DNA) adapted for 96-well format and automated on the Labcyte Echo 525 and Lynx by Dynamic Devices LM1200 liquid handler. A 1 in 10 dilution of each transformation mix was plated as a 5 μL droplet on LB agar + kanamycin (50 µg/mL) and incubated at 37 °C overnight. A single colony of each clone was picked into 2 mL 96-well V-bottom deep-well plates (Azenta) containing 1 mL of Terrific Broth (TB) + 50 μg/mL kanamycin using a Singer PIXL automated colony picker, and grown overnight at 37 °C, 600 rpm. All 96-well plate cultures were grown in an Infors HT multitron with 0.3 cm throw. Strains containing pET29b-eGFP, pET29b-ppEST and pET29b-N169 (unpublished PpEST-family ancestral protein) were included on each culture plate as negative and positive controls, respectively. Cultures were normalised to OD₆₀₀ = 0.1 on the Lynx LM900 96VVP liquid handler with on deck Byonoy Absorbance 96, and induced with 1 mM IPTG at 37 °C, 600 rpm. OD_600_ was recorded after induction. After induction for 3 h, cells were pelleted (4,000 rpm, 10 min) and stored at −80 °C until lysis.

#### High-throughput esterase activity assays

Cell pellets were lysed in NEBExpress® E. coli Lysis Reagent (New England Biolabs) supplemented with 0.1 mg /mL lysozyme (Merck), Benzonase (Merck), and protease inhibitor cocktail (for Poly-His proteins, P 8849, Merck) according to manufacturer’s instructions. Lysis was performed in 96-well plates (100 µL/well) at 20 °C, 900 rpm for 20 min. Lysates were clarified by centrifugation (4,000 rpm, 10 min, 4 °C) and diluted 1:10 in 50 mM Tris buffer (pH 8.0). Diluted lysates were transferred to 384-well Echo PP plates (Beckman) for dispensing on the Labcyte Echo 525.

Esterase activity was measured against the *p*-nitrophenyl (pNP) esters: pNP-acetate (C2), pNP-butyrate (C4), and pNP-octanoate (C8). Stocks were made at 15 mM in methanol, prepared fresh and stored at −20 °C. Reactions were assembled in 96-well flat-bottom plates (Corning) by dispensing 2 µL lysate in triplicate and 93 µL of 50 mM Tris buffer pH 8.0. For thermostability screening, lysates were first incubated at 60 °C and 90 °C for 10 min in 384-well PCR plates (Azenta), cooled to 2 °C, and then transferred to Echo LDV plates for assay setup. To run the assay, 5 µL of substrate was added to plates using the injector on a BioteK Cytation 5 plate reader, for a final substrate concentration of 750 µM in 5 % methanol. Hydrolysis was monitored by measuring absorbance or optical density (OD) at 410 nm at 24 °C for 2 min 30 sec (read every 11 sec). Three blank wells containing only buffer and substrate in 100 μl total volume were included on each assay plate. The data reduction method “blank transformation” on Cytation Gen5 software (version 3.15.15) was set up to automatically subtract the average of the blank reads from all the OD values before calculating the MaxV (mOD/min). This is the calculated value of the mean slope from a linear regression of points in a calculation zone. In this case, the calculation zone was set to include five time points. Maximum velocity (MaxV) was used as the comparative metric across the dataset. The software also reported R-squared values, which were used for quality control. Data points were excluded from the data set if the R^2^ was < 0.96 (rejected data).

The esterase from porcine liver (Merck: E3019; CAS Number: 9016-18-6) and the PpEST-family ancestral protein N169 (unpublished) were included in triplicate at approximately 0.003 mg/mL and 0.02 μM final concentration in 50 mM Tris pH 8.0, respectively, as controls on each assay plate. The assays were repeated if the results for these controls were inconsistent.

Strains were tracked from transformation through to the assays using Teselagen (AI-Assisted Software for Life Sciences - Teselagen).

### Pretrained and physics-based baselines

To evaluate the suitability of existing computational approaches for enzyme property prediction, we applied molecular docking (GNINA^44^) and eight pre-trained models (CLIPZyme^45^, ProSmith^46^, CatPred^47^, SELFprot^48^, DeepSTABp^49^, TemBERTure^50^, NetSolP^51^, PLM_Sol^52^) to predict properties for all 2,123 PpEST-like sequences.

Pre-trained machine learning models were used to estimate enzyme substrate compatibility, catalytic efficiency, thermostability, and solubility. Using pNP-acetate, pNP-butyrate or pNP-octanoate as substrates, four models were used to predict substrate-specific properties: CLIPZyme ^45^ to predict enzyme-reaction probability, ProSmith^46^ to predict enzyme-substrate probability, and CatPred^47^ and SELFProt^48^ to each predict both k_cat_ and K_M_. DeepSTABp^49^ and TemBERTure^50^ were used to predict the melting temperature of each enzyme. PLM_Sol^52^ and NetSolP^51^ were used for solubility predictions.

To obtain input structures for molecular docking, protein structures were predicted using AlphaFold3^37^ and conformers were generated for the three substrates: pNP-acetate (five conformers per isomer), pNP-butanoate (10 conformers per isomer), and pNP-octanoate (20 conformers per isomer), using RDkit. Covalent docking was performed with GNINA v1.1.0 software on all conformers of both stereoisomers of the acyl-enzyme intermediate state, where the carbonyl carbon of the substrate is covalently bound to the catalytic serine. The catalytic serine was identified for each structure by superposing all the structures and selecting the serine oxygen within 2 Å of the identified catalytic serine from sequence conservation. For each enzyme-substrate pair, three docking scores were calculated via GNINA^44^: vina affinity, CNN affinity and CNN VS (CNN affinity * CNN score). Exhaustiveness was set to 64. The average of the top seven scores were calculated for each stereoisomer and the best score was used for downstream analysis with experimental results.

For overall activity, correlations were first evaluated for each predictor independently. Only two predictors CatPred k_cat_ (ρ ≈ 0.19) and NetSolP solubility (ρ ≈ 0.21) had correlations above 0.1. Rank aggregation of the two predictors resulted in an improved correlation (ρ ≈ 0.28). For octanoate preference, correlations were assessed separately for each substrate. Similar to overall activity, only CatPred k_cat_ and NetSolP models had ρ > 0.1 for all three substrates. Notably, both models showed stronger correlations for octanoate than acetate: CatPred k_cat_ (ρ ≈ 0.22 vs. 0.10) and NetSolP (ρ ≈ 0.33 vs. 0.14). GNINA CNN VS scores were also positively correlated when filtered for octanoate (ρ ≈ 0.11), whereas acetate scores were inversely correlated (ρ ≈ -0.18). For this study, as octanoate preference is defined as the log ratio of octanoate to acetate activity, only octanoate-specific scores were retained for the final prediction. Aggregating CatPred k_cat_, NetSolP and GNINA CNN VS results increased the correlation to ρ ≈ 0.41. For thermostability, we used only models are trained on melting temperature datasets (DeepSTABp and TemBERTure). DeepSTABp was more strongly correlated to measured thermostability (ρ ≈ 0.14) compared to TemBERTure (ρ ≈ 0.07). This procedure simulates a realistic use-case in which heterogeneous computational tools are combined to prioritise select sequences in the absence of supervised training.

Spearman’s correlations were used to compare molecular docking and pre-trained model scores with experimental results. To process experimental results for analysis, overall activity, thermostability and substrate specificity were calculated for each sequence as defined in the previous section. For overall activity, correlations were first calculated separately for each docking and pre-trained model score. Methods with an initial correlation greater than 0.1 were combined and used to re-rank sequences with the rank aggregation procedure^53^. A final correlation was then computed between the re-ranked sequences and overall activity ranks obtained from experimental results. For thermostability, correlations were computed individually for the DeepSTABp^49^ and TemBERTure^50^ models only. For substrate specificity, analysis was performed in three stages. First, correlations were calculated between measured maxV and model scores for each substrate separately. As octanoate correlations were stronger than other substrates, CLIPZyme^45^, ProSmith^46^, CatPred^47^ and SELFprot^48^ scores were filtered for octanoate-specific predictions. Next, correlations were calculated between each model and measured octanoate specificity. Sequences were re-ranked with the rank aggregation procedure using models with a correlation greater than 0.1. Then, to check that the combined scores improved predictions, a final correlation was calculated for the aggregated ranks.

### Family-specific machine learning

#### Target construction

For constructing a dataset to train supervised machine learning models, we denote experimental MaxV values as 𝑦_V-Acetate_, 𝑦_V-Butyrate_, 𝑦_V-Octanoate_, 𝑦_V-Butyrate-60_, 𝑦_V-Butyrate-90_. We use these values to derive labels representing our objectives; overall activity, thermostability and substrate specificity. We clamped the MaxV’s to have a minimum value of 1 to avoid singular ratios in the derived labels. The derived labels are defined as:

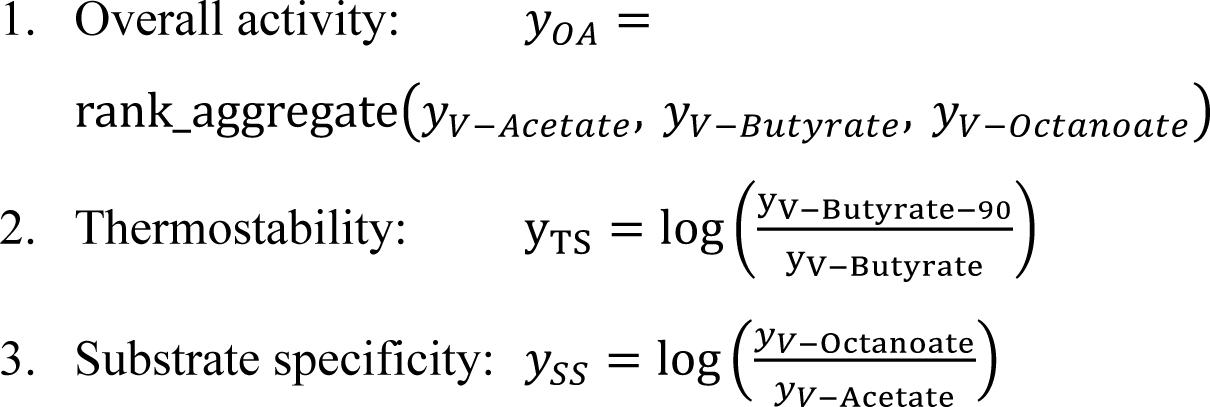

Only sequences with denominator values 𝑦 > 10 were included in (2) and (3) to prevent low-activity samples from dominating the ratios. The rank aggregation procedure follows Bedo *et al.* (2016) ^53^: MaxV values for acetate, butyrate and octanoate substrates were ranked 𝑠𝑡𝑎𝑟𝑡𝑖𝑛𝑔 𝑎𝑡 𝑟_*t*_ = 1 for the highest value, with increasing integers indicating lower ranks. These ranks were then normalised by 𝑟̃_*t*_ = 𝑟_*t*_/max (𝑟_*t*_ + 1), and finally combined geometrically as 𝑦_OA_ = 1 − exp [(1/3) ∑_*t*_ log 𝑟_*t*_], yielding an aggregate score between 0 (worst) and 1 (best). This procedure maximises the spearman correlation between the final ranking score, 𝑦_OA_, and the original ranks of each substrate MaxV.

#### Model architecture

Initially we trialled three neural network architectures for predicting the preceding labels: (1) A two layer convolutional neural network (CNN)^54^ on embedded, integer-encoded (token) sequences with end zero-padding, (2) a CNN on integer-encoded multiple sequence alignment (MSA) inputs with the same topology and (3) a transformer encoder applied to MSA sequences. Using 5-fold cross validation on the 1518 sequence dataset we found that architecture (1) outperformed the others for all labels, and was used for all subsequent work. The base CNN architecture is depicted in **Figure S13** panel B and is adapted from Steinberg *et al.* (2025).^55^

#### Training and validation

The neural network parameters were optimised by minimising the mean squared error (MSE), or the weighted MSE (WMSE) of the model predictions depending on the selection method used (detailed in the next section). A small 𝑙_3_ penalty was applied to the network weights, with the strength of this penalty chosen using cross validation. All models were trained for 50 epochs with AdamW^56^.

Model generalisation was quantified using uniform random five-fold cross-validation^57^ with uniform random sampling of the original 1518 sequences and corresponding targets (no stratified or grouped sampling strategy was required). Performance metrics included out-of-fold Spearman’s ρ and R² (**Table 1**). Spearman correlation is meaningful in our setting as we are interested in the highest ranked enzymes for further analysis, and not the precise value of MaxV obtained.

After cross validation, the models were trained from scratch on all 1,518 training sequences and targets for 50 epochs. These models were then used for selecting highest ranked candidates from the remaining 605 untested sequences.

#### Data selection strategies

We trialled two selection strategies using the predictive models: highest expected value (HEV) and upper confidence bound (UCB).

#### Highest expected value

(HEV) – this selection strategy (referred to as “predictive mean” in Garnet (2023)^35^) simply entails selecting the sequences with the highest predicted target values from the pool of 605 sequences. The predictor used for selection is a single CNN trained using WMSE. Normalised inverse (discretised) frequency weights were used for WMSE such that sequences with less frequently occurring targets were upweighted. This has the effect of making the predictor equally concerned with fitting rarer sequences with high and low valued targets as those more common middle-valued. This disincentivises “regressing to the mean” as is typical with unweighted MSE.

#### Upper confidence bound

(UCB) – this strategy entails selecting the sequences with the highest predicted 90^th^-percentile predictive-interval^35^. This strategy attempts to incorporate model (epistemic) uncertainty into the selection criterion, and gives sequences that have a less confident prediction an “exploration bonus” as opposed to the purely exploitative HEV selection criterion above. As such this can be viewed as optimistic strategy with respect to the uncertainty of our predictions. For this we need a predictive distribution, rather than just the predicted expected value required by HEV. We approximate predictive distribution by training an ensemble of 10 separate randomly-initialised CNNs using MSE, and then sampling predictions from them using test-time enable weight dropout^58^. We gather 100 samples (10 from each CNN in the ensemble) to construct an empirical predictive 90^th^-percentile value for each of the 605 sequences. In this case we train the CNNs using MSE (using the average of their individual predictions as the predicted expected value) since we want to preserve the calibration of the predictive density for estimating percentiles.

### Candidate panel generation

Predictions for 605 untested *SGNH* sequences were ranked for each task using three strategies (CNN-HEV, CNN-UCB, and generalist pre-trained model prediction). To establish a baseline, the 605 untested sequences were clustered using CD-HIT^38^ at a 46% identity threshold (c = 0.46), yielding 30 representative sequences. This baseline was then merged with the top-K predictions from the nine ranked lists. Unique entries were retained while incrementally increasing K until the deduplicated union reached the experimental budget of 230 sequences at K = 50. This yielded a diverse and balanced selection for prospective validation.

### Residue predictive importance

To interpret which sequence positions most strongly influenced model predictions, we applied Integrated Gradients (IG)^21^ attribution to the CNN ensemble predictor. Integrated Gradients is a gradient-based method that assigns an importance score to each input position by measuring how changes in the input representation affect the model’s prediction. This provides a residue-level estimate of how strongly each amino-acid position contributes to the predicted biochemical property.

Attributions were computed with respect to the embedded sequence representation provided to the first layer of the CNN. In our model, integer-encoded amino-acid sequences are first mapped to dense continuous embeddings before being processed by the convolutional layers. Integrated Gradients was therefore applied to these embedding vectors rather than the discrete token indices, allowing the method to quantify the contribution of each residue position in a smooth and well-defined manner.

As a reference input (baseline), we used an all-padding sequence composed entirely of the padding token (“–”), corresponding to the zero-padded representation used during model training. This baseline represents a sequence lacking informative amino-acid content, enabling the attribution scores to capture the incremental contribution of each observed residue relative to an absence of sequence signal.

For each sequence and prediction task, attribution scores were computed for the CNN ensemble and then aggregated to produce a single per-residue importance profile. Scores were obtained by combining the signed contributions across embedding dimensions at each sequence position and subsequently normalised for visualisation. These per-position importance values were mapped onto multiple sequence alignments and representative protein structures to highlight regions where sequence variation most strongly influenced predicted activity, substrate specificity, and thermostability.

This procedure provides a task-specific measure of how local sequence patterns contribute to model predictions, while remaining consistent with the sequence-only nature of the predictive framework used throughout this study.

### Benchmarking selection efficiency via simulation

To compare the efficiency of different selection strategies, we performed an offline simulation using the assayed sequences from the first and second experimental cohorts (1,743 sequences) as a ground-truth domain. The simulation was designed to mimic the iterative cycle of a real-world sampling campaign. To ensure a fair baseline, selection strategies were compared using the same set of diverse starting sequences generated using CD-HIT^38^ at 40% sequence identity. The simulation proceeded in batched rounds where each selection strategy was able to nominate 10 unobserved sequences to be revealed by the ground-truth oracle, mimicking an iterative experimental sampling campaign. While physical high-throughput workflows utilise standard plate formats (e.g. 96-well), simulating a smaller batch size provides a higher-resolution assessment of the learning dynamics and establishes the theoretical discovery potential of adaptive strategies over more passive methods.

Four strategies were evaluated: random sampling, pretrained models, CNN-HEV, and a CNN-UCB ensemble. The random selection strategy is a non-adaptive baseline that draws a random batch from the remaining pool in each iteration. The pretrained model strategy is a deterministic and non-adaptive strategy that only depend on the pool of input sequences. It follows a fixed ranking of sequences and does not update with new data. As such, the strategy iteratively works down a fixed ranking of the sequence pool in batches. The two CNN strategies employ an adaptive, model-in-the-loop approach where they are retrained on all previously observed data at every iteration. This enables the models to re-rank the entire remaining pool of sequences before each batch selection. To account for stochastic variation, the non-deterministic strategies, random selection and both CNN strategies, were evaluated across 10 independent runs with different random seeds. Note that since the pretrained model strategy is deterministic, it was only run once.

Performance was assessed using three complementary metrics that represent distinct priorities in a protein engineering campaign: cumulative hit rate, recall, and cumulative regret. Hits were defined as sequences whose measured performance for a given task fell within the top 10% (90th percentile) of the complete dataset. Cumulative hit rate (or yield) is non-monotonic and quantifies sampling efficiency by calculating the fraction of selected sequences that qualify as hits. A highly effective strategy will show an early peak as it successfully identifies high performers, followed by a gradual decline as the finite pool of hits is exhausted. Recall (or sensitivity) measures the breadth of discovery by tracking the proportion of total available hits identified within a given sampling budget. This metric is monotonically increasing, as discovery coverage can only increase or plateau as more sequences are observed. Finally, cumulative regret quantifies the opportunity cost of a selection strategy by summing the difference in performance between the observed batch and the best possible batch available at each iteration. A lower cumulative regret indicates a strategy that consistently prioritises top-tier variants, effectively minimising the shortfall relative to an optimal sampling trajectory.

## Data availability

Data generated during the study are available as supplementary information files.

## Code availability

Code used in this work are available in supplementary information files.

## Acknowledgements

This project was supported by resources and expertise provided by CSIRO IMT Scientific Computing.

## Author contributions

**Conceptualisation, design and analysis:** All Authors. **Writing, review & editing:** All Authors. **Methodology:** F.H.A. (structural biology, protein evolution, biochemistry), D.S. (machine learning, feature importance), A.B. (machine learning, simulation/benchmarking), H.P., A.C.W (pretrained models)., L.G. and C.J. (laboratory experiments). **Software and data management:** A.B., D.S., A.M. **Laboratory experiments:** F.H.A., L.G., C.I., W.H., C.J. (high-throughput screening and experimental characterisation). **Project administration and supervision:** C.S.O., R.E.S.

## References

1. Moorhoff, F. et al. Machine Learning-Driven Enzyme Mining: Opportunities, Challenges, and Future Perspectives. ACS Catal. 16, 12–30 (2025).

2. Du, Q., Wang, H., Jiang, B. & Wang, X. Advancing genetic engineering with active learning: theory, implementations and potential opportunities. Brief. Bioinform. 26, 286 (2025).

3. Ao, Y. F. et al. Data-Driven Protein Engineering for Improving Catalytic Activity and Selectivity. ChemBioChem 25, e202300754 (2024).

4. Hie, B. L. & Yang, K. K. Adaptive machine learning for protein engineering. Curr. Opin. Struct. Biol. 72, 145–152 (2022).

5. Romero, P. A. & Arnold, F. H. Exploring protein fitness landscapes by directed evolution. Nat. Rev. Mol. Cell Biol. 10, 866–876 (2009).

6. Muir, D. F. et al. Evolutionary-scale enzymology enables exploration of a rugged catalytic landscape. Science (1979). 388, (2025).

7. Notin, P., Rollins, N., Gal, Y., Sander, C. & Marks, D. Machine learning for functional protein design. Nature Biotechnology 2024 42:2 42, 216–228 (2024).

8. Paton, A. E. et al. Connecting chemical and protein sequence space to predict biocatalytic reactions. Nature 2025 646:8083 646, 108–116 (2025).

9. Bileschi, M. L. et al. Using deep learning to annotate the protein universe. Nature Biotechnology 2022 40:6 40, 932–937 (2022).

10. Capela, J. et al. Comparative Assessment of Protein Large Language Models for Enzyme Commission Number Prediction. BMC Bioinformatics 26, (2025).

11. Schmirler, R., Heinzinger, M. & Rost, B. Fine-tuning protein language models boosts predictions across diverse tasks. Nature Communications 2024 15:1 15, 7407- (2024).

12. Gruver, N. et al. Protein Design with Guided Discrete Diffusion. Adv. Neural Inf. Process. Syst. 36, (2023).

13. Jumper, J. et al. Highly accurate protein structure prediction with AlphaFold. Nature 2021 596:7873 596, 583–589 (2021).

14. Rao, R. et al. Evaluating Protein Transfer Learning with TAPE. Adv. Neural Inf. Process. Syst. 32, 9689 (2019).

15. Dallago, C. et al. FLIP: Benchmark tasks in fitness landscape inference for proteins. *bioRxiv* Preprint at https://www.biorxiv.org/content/10.1101/2021.11.09.467890v1 (2021).

16. Freschlin, C. R., Fahlberg, S. A., Heinzelman, P. & Romero, P. A. Neural network extrapolation to distant regions of the protein fitness landscape. Nature Communications 2024 15:1 15, 6405- (2024).

17. Matthews, D. S. et al. Leveraging ancestral sequence reconstruction for protein representation learning. Nature Machine Intelligence 2024 6:12 6, 1542–1555 (2024).

18. Norton-Baker, B. et al. Machine Learning-Guided Identification of PET Hydrolases from Natural Diversity. ACS Catal. 15, 16070–16083 (2025).

19. Goldman, S., Das, R., Yang, K. K. & Coley, C. W. Machine learning modeling of family wide enzyme-substrate specificity screens. PLoS Comput. Biol. 18, e1009853 (2022).

20. Yang, J. et al. Active learning-assisted directed evolution. Nature Communications 2025 16:1 16, 714- (2025).

21. Axiomatic attribution for deep networks | Proceedings of the 34th International Conference on Machine Learning - Volume 70. https://dl.acm.org/doi/10.5555/3305890.3306024.

22. Paysan-Lafosse, T. et al. The Pfam protein families database: embracing AI/ML. Nucleic Acids Res. 53, D523–D534 (2025).

23. Bateman, A. et al. UniProt: the Universal Protein Knowledgebase in 2025. Nucleic Acids Res. 53, D609–D617 (2025).

24. Blum, M. et al. The InterPro protein families and domains database: 20 years on. Nucleic Acids Res. 49, D344–D354 (2021).

25. Clifton, B. E., Kozome, D. & Laurino, P. Efficient Exploration of Sequence Space by Sequence-Guided Protein Engineering and Design. Biochemistry 62, 210–220 (2022).

26. Gerlt, J. A. et al. Enzyme Function Initiative-Enzyme Similarity Tool (EFI-EST): A web tool for generating protein sequence similarity networks. Biochim. Biophys. Acta 1854, 1019–37 (2015).

27. Akram, F. et al. Abridgement of Microbial Esterases and Their Eminent Industrial Endeavors. Molecular Biotechnology 2024 67:3 67, 817–833 (2024).

28. Anderson, A. C., Stangherlin, S., Pimentel, K. N., Weadge, J. T. & Clarke, A. J. The SGNH hydrolase family: a template for carbohydrate diversity. Glycobiology 32, 826–848 (2022).

29. Denessiouk, K. et al. The active site of the SGNH hydrolase-like fold proteins: Nucleophile–oxyanion (Nuc-Oxy) and Acid–Base zones. Curr. Res. Struct. Biol. 7, 100123 (2024).

30. Wallace, P. W. et al. PpEst is a novel PBAT degrading polyesterase identified by proteomic screening of Pseudomonas pseudoalcaligenes. Appl. Microbiol. Biotechnol. 101, 2291–2303 (2017).

31. Ahmed, F. H. et al. Disordered N-termini enhance the thermostability of SGNH-hydrolase family polyesterases. Protein Science 35, e70402 (2026).

32. Li, Z. et al. Structure-guided protein engineering increases enzymatic activities of the SGNH family esterases. Biotechnol. Biofuels 13, (2020).

33. Lin, Z. et al. Evolutionary-scale prediction of atomic-level protein structure with a language model. Science (1979). 379, 1123–1130 (2023).

34. Ahmed, F. H. et al. Thermostable Bacterial Esterases From Lipase Family 1.5 Degrade Compostable Polyesters PBAT and PBSA. Microbiologyopen 14, e70144 (2025).

35. Garnett, R. Bayesian Optimization. Bayesian Optimization (Cambridge University Press, 2023).

36. Procter, J. B. et al. Alignment of Biological Sequences with Jalview. Methods Mol. Biol. 2231, 203 (2021).

37. Abramson, J. et al. Accurate structure prediction of biomolecular interactions with AlphaFold 3. Nature 2024 630:8016 630, 493–500 (2024).

38. Fu, L., Niu, B., Zhu, Z., Wu, S. & Li, W. CD-HIT: accelerated for clustering the next-generation sequencing data. Bioinformatics 28, 3150–3152 (2012).

39. Teufel, F. et al. SignalP 6.0 predicts all five types of signal peptides using protein language models. Nature Biotechnology 2022 40:7 40, 1023–1025 (2022).

40. Sainburg, T., McInnes, L. & Gentner, T. Q. Parametric UMAP Embeddings for Representation and Semi-supervised Learning. Neural Comput. 33, 2881 (2021).

41. Rozewicki, J., Li, S., Amada, K. M., Standley, D. M. & Katoh, K. MAFFT-DASH: integrated protein sequence and structural alignment. Nucleic Acids Res. 47, W5–W10 (2019).

42. Minh, B. Q. et al. IQ-TREE 2: New Models and Efficient Methods for Phylogenetic Inference in the Genomic Era. Mol. Biol. Evol. 37, 1530–1534 (2020).

43. Letunic, I. & Bork, P. Interactive Tree of Life (iTOL) v6: recent updates to the phylogenetic tree display and annotation tool. Nucleic Acids Res. 52, W78–W82 (2024).

44. McNutt, A. T. et al. GNINA 1.0: molecular docking with deep learning. Journal of Cheminformatics 2021 13:1 13, 43- (2021).

45. Mikhael, P. G., Chinn, I. & Barzilay, R. CLIPZyme: Reaction-Conditioned Virtual Screening of Enzymes. Proc. Mach. Learn. Res. 235, 35647–35663 (2024).

46. Kroll, A., Ranjan, S. & Lercher, M. J. A multimodal Transformer Network for protein-small molecule interactions enhances predictions of kinase inhibition and enzyme-substrate relationships. PLoS Comput. Biol. 20, e1012100 (2024).

47. Boorla, V. S. & Maranas, C. D. CatPred: a comprehensive framework for deep learning in vitro enzyme kinetic parameters. Nature Communications 2025 16:1 16, 2072-(2025).

48. Wilson, M., Coudrat, T. & Warden, A. SELFprot: Effective and Efficient Multitask Finetuning Methods for Protein Parameter Prediction. J. Chem. Inf. Model. 65, 3226–3238 (2025).

49. Jung, F., Frey, K., Zimmer, D. & Mühlhaus, T. DeepSTABp: A Deep Learning Approach for the Prediction of Thermal Protein Stability. International Journal of Molecular Sciences 2023, Vol. 24, 24, (2023).

50. Rodella, C., Lazaridi, S. & Lemmin, T. TemBERTure: advancing protein thermostability prediction with deep learning and attention mechanisms. Bioinformatics Advances 4, (2024).

51. Thumuluri, V. et al. NetSolP: predicting protein solubility in Escherichia coli using language models. Bioinformatics 38, 941–946 (2022).

52. Zhang, X. et al. PLM_Sol: predicting protein solubility by benchmarking multiple protein language models with the updated Escherichia coli protein solubility dataset. Brief. Bioinform. 25, (2024).

53. Bedö, J. & Soon, O. Multivariate spearman’s ρ for aggregating ranks using copulas. The Journal of Machine Learning Research Preprint at https://dl.acm.org/doi/pdf/10.5555/2946645.3053483 (2016).

54. Li, Z., Liu, F., Yang, W., Peng, S. & Zhou, J. A Survey of Convolutional Neural Networks: Analysis, Applications, and Prospects. IEEE Trans. Neural Netw. Learn. Syst. 33, 6999–7019 (2022).

55. Steinberg, D. M., Oliveira, R., Ong, C. S. & Bonilla, E. V. Variational Search Distributions. 13th International Conference on Learning Representations, ICLR 2025 2812–2851 (2025).

56. Loshchilov, I. & Hutter, F. Decoupled Weight Decay Regularization. 7th International Conference on Learning Representations, ICLR 2019 http://arxiv.org/abs/1711.05101 (2019).

57. Hastie, T., Tibshirani, R. & Friedman, J. *The Elements of Statistical Learning*. (Springer New York, New York, NY, 2009).

58. Gal, Y. & Ghahramani, Z. Dropout as a Bayesian Approximation: Representing Model Uncertainty in Deep Learning. 1050–1059 Preprint at https://proceedings.mlr.press/v48/gal16.html (2016).

