## Supplementary Figures for "Data-Efficient Exploration of Enzyme Function Using Family-Specific Machine Learning"

OFFICIAL

OFFICIAL

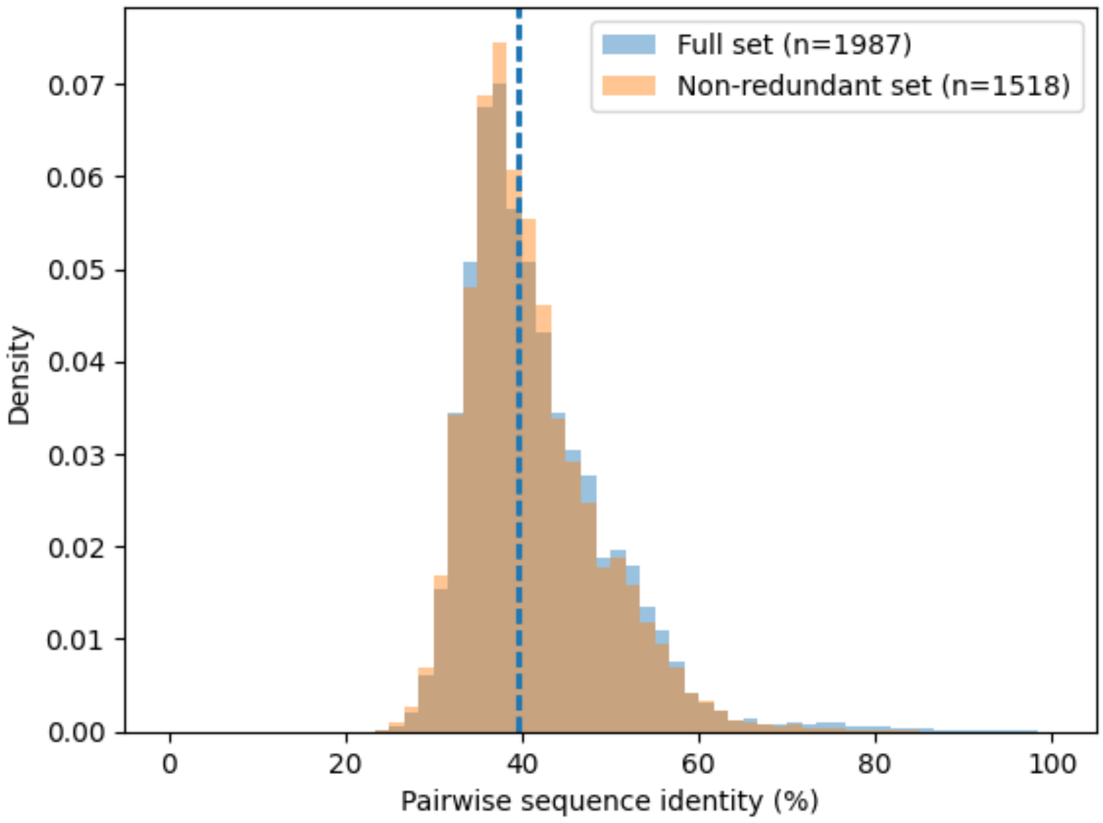


**Figure S1. Pairwise sequence identity distribution within the PpEST clade characterised in this work.** Pairwise sequence identities were calculated from the multiple sequence alignment for the full dataset excluding the related sister clade (n = 1,987 sequences) and the non-redundant dataset obtained after applying an 89.5% identity filter using CD-HIT[1] (n = 1,518 sequences). Histograms show normalised density from sampled pairwise comparisons (250,000 pairs per dataset). The median is shown as a dotted line.

**Table S1. Automated biofoundry workflow used for high-throughput screening of SGNH-hydrolase homologues.** The workflow integrates acoustic liquid handling, automated colony picking, standardised expression, chemical lysis, thermostability treatment, and kinetic activity measurements using p-nitrophenyl substrates. Quality control checkpoints were implemented at each stage to ensure reproducible enzyme expression and activity measurements across plates.

| **Step** | **Platform / Instrument** | **Process** | **Quality control** |
|---|---|---|---|
| Miniaturised transformation | Echo 525 acoustic liquid handler; Lynx LM1200 liquid handler | Heat-shock transformation of codon-optimised pET-29b constructs into *E. coli* T7 Express lysY cells in 96-well format | Transformation conditions validated across DNA input concentrations and dispensing volumes |
| Colony picking | Singer PIXL automated colony picker | Single colonies transferred into 96-well deep-well expression plates | Automated colony selection and plate layout verification |
| Expression culture | Infors HT Multitron shaker incubator, 3mm throw. | Cultures grown in TB medium and induced with IPTG | Control strains included on each plate |
| Culture normalisation | Lynx LM900 liquid handler with Byonoy Absorbance 96 | $OD_{600}$ measured and cultures normalised prior to lysis | Automated blank subtraction and OD calculation |
| Cell lysis | Lynx LM1200 liquid handler; Infors HT Multitron shaker | Chemical lysis and clarification of cell pellets | Lysis volumes and dilution conditions validated |
| Thermostability treatment | Lynx LM1200 liquid handler | Lysates incubated at elevated temperatures prior to assay | Reproducibility of heat treatment verified across plates |
| Assay setup | Lynx liquid handlers; Echo 525 | Automated dispensing of lysate and pNP substrates into assay plates | Liquid handling volumes validated |
| Esterase activity assay | BioTek Cytation 5 plate reader | Hydrolysis of pNP substrates monitored kinetically at 410 nm | Assays performed in triplicate with blank wells |
| Data processing | Gen5 software | MaxV calculated from linear regression of kinetic traces | Data filtered using $R^2$ threshold and plate controls |
| Plate controls | BioTek Cytation 5 plate reader | Reference enzymes included on every plate | Assays repeated if control activity deviated |
| Sample tracking | Teselagen LIMS (https://teselagen.com/platform) | Construct identity tracked from transformation to assay | Digital tracking of all constructs and assays |

OFFICIAL

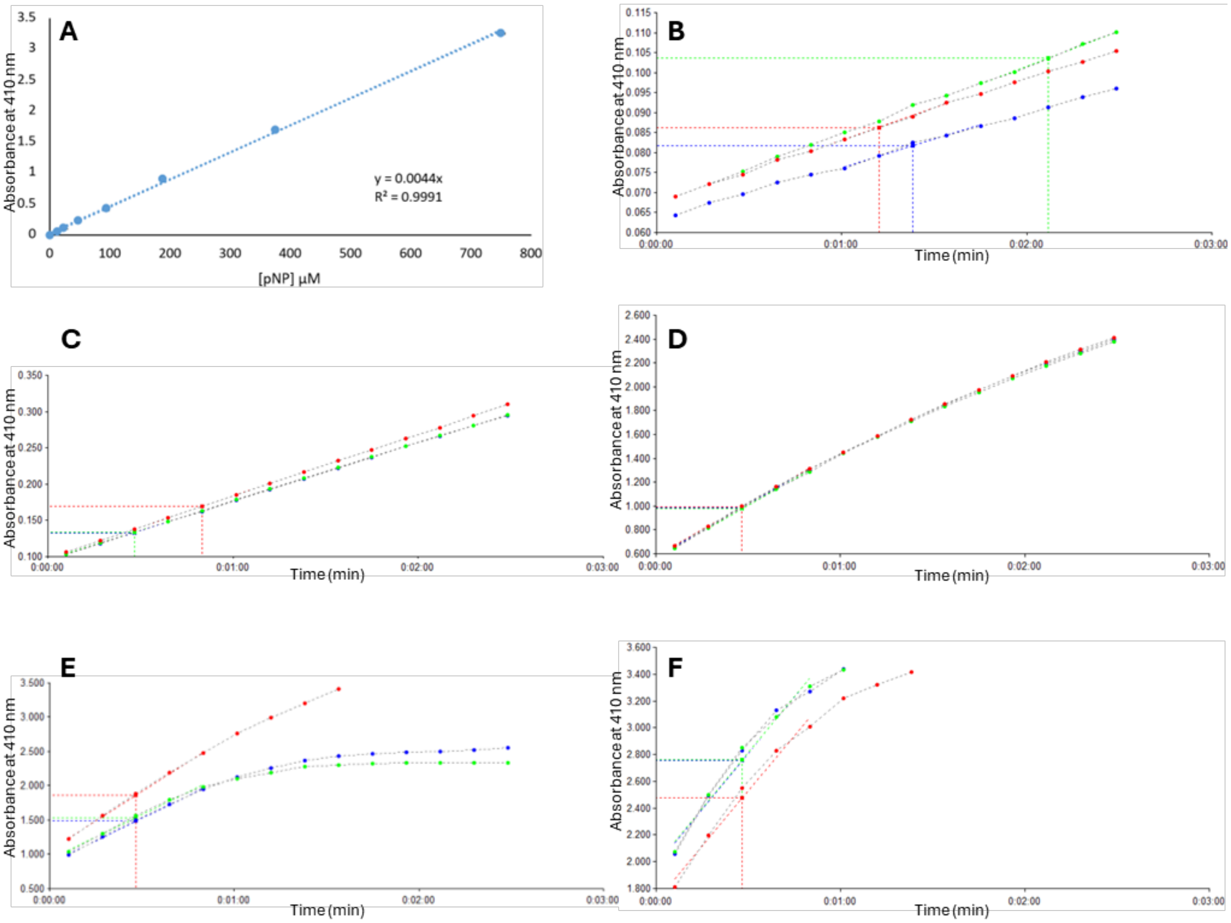


**Figure S2. Validation of the pNP esterase activity assay used for high-throughput screening. A.** Calibration curve for *p*-nitrophenol measured at 410 nm following serial dilution in assay buffer. Absorbance increased linearly with concentration across the detection range used for activity measurements ($R^2 = 0.999$). **B-E.** Representative traces illustrating the range of activities observed during screening. Absorbance at 410 nm was monitored following addition of pNP-butyrate, and reaction velocities were estimated from the linear phase of absorbance increase using automated regression in the Cytation Gen5 analysis software. **B.** Background signal from a blank reaction, **C.** Low activity enzyme, **D.** Moderate activity enzyme, and **E.** High activity enzyme. **F.** Example of detector saturation at very high activity, illustrating the upper detection limits of the assay. Reaction velocities used in downstream analyses were calculated only from the linear region prior to saturation.

OFFICIAL

OFFICIAL

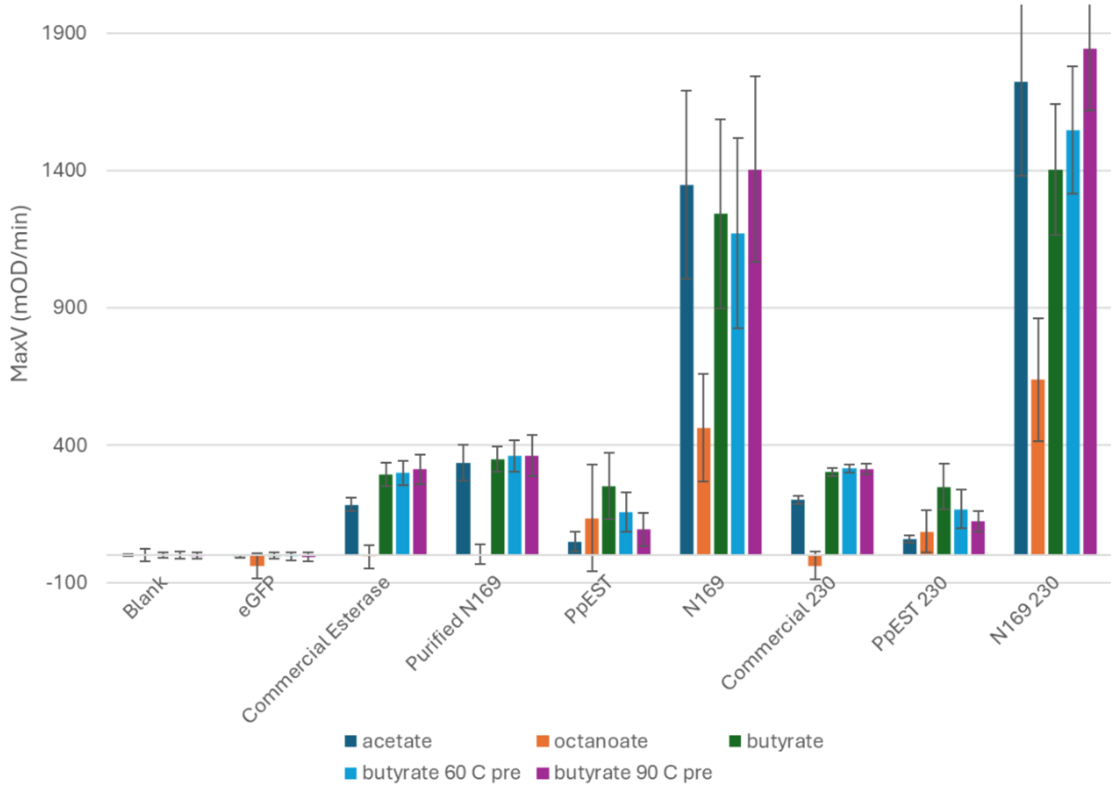


**Figure S3. Performance of assay controls across screening and validation campaigns.** Blank wells, eGFP-negative controls, and reference enzymes were included throughout both the initial high-throughput screen (1513 sequences) and the second experimental batch used to validate the 230 model-selected sequences. Reference enzymes included a commercial porcine liver esterase, the previously characterised PpEST enzyme, and the reconstructed ancestral SGNH-hydrolase N169 (unpublished), which served as internal activity standards. Bars show mean maximum velocity (MaxV, mOD/min) values measured for each enzyme under the assay conditions tested: pNP-acetate, pNP-octanoate, pNP-butyrate, and pNP-butyrate following pre-incubation at 60 °C or 90 °C. Controls labelled "230" correspond to the second experimental batch used for prospective validation of the machine-learning selected sequences. Error bars represent the standard deviation across replicate measurements: Commercial esterase and purified N169 in 1513 assay experiments n = 207; Commercial esterase and purified N169 in 230 assay experiments n = 27; PpEST and N169 lysate in 1513 assays n = approximately 50 (varied slightly for each assay after data quality control); PpEST and N169 lysate in 230 assays n = 9.

OFFICIAL

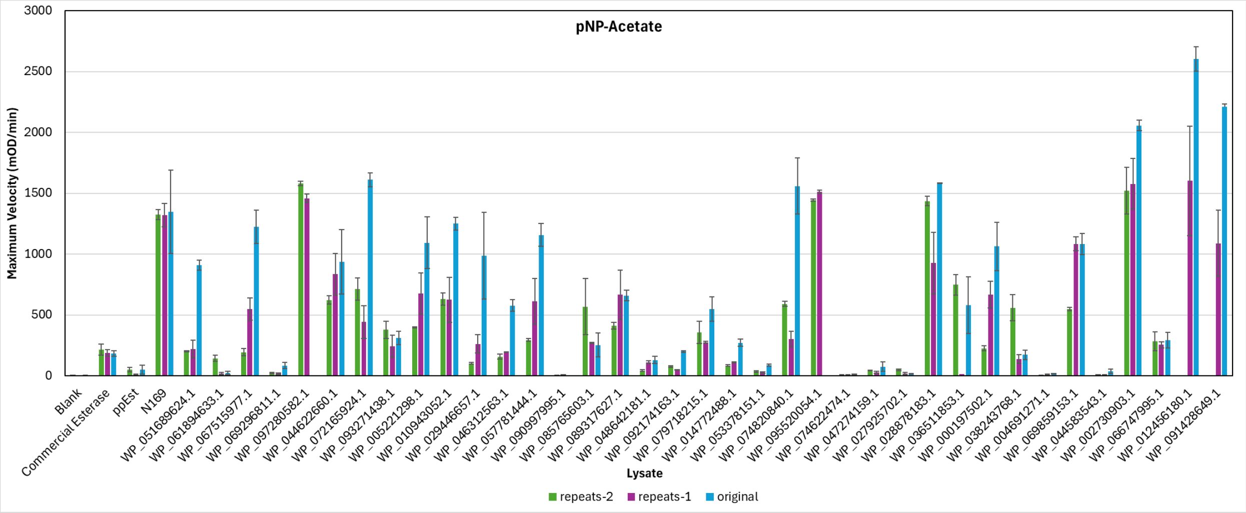
pNP-Acetate
Maximum Velocity (mOD/min)
3000
2500
2000
1500
1000
500
0
Blank
Commercial Esterase
ppEst
N169
WP_051689624.1
WP_061894633.1
WP_067515977.1
WP_069296811.1
WP_097280582.1
WP_044622660.1
WP_072165924.1
WP_093271438.1
WP_005221298.1
WP_010943052.1
WP_029446657.1
WP_046312563.1
WP_057781444.1
WP_090997995.1
WP_085765603.1
WP_089317627.1
WP_048642181.1
WP_092174163.1
WP_079718215.1
WP_014772488.1
WP_053378151.1
WP_074820840.1
WP_095520054.1
WP_074622474.1
WP_047274159.1
WP_027925702.1
WP_028878183.1
WP_036511853.1
WP_000197502.1
WP_038243768.1
WP_004691271.1
WP_069859153.1
WP_044583543.1
WP_002730903.1
WP_066747995.1
WP_012456180.1
WP_091428649.1
Lysate
repeats-2
repeats-1
original

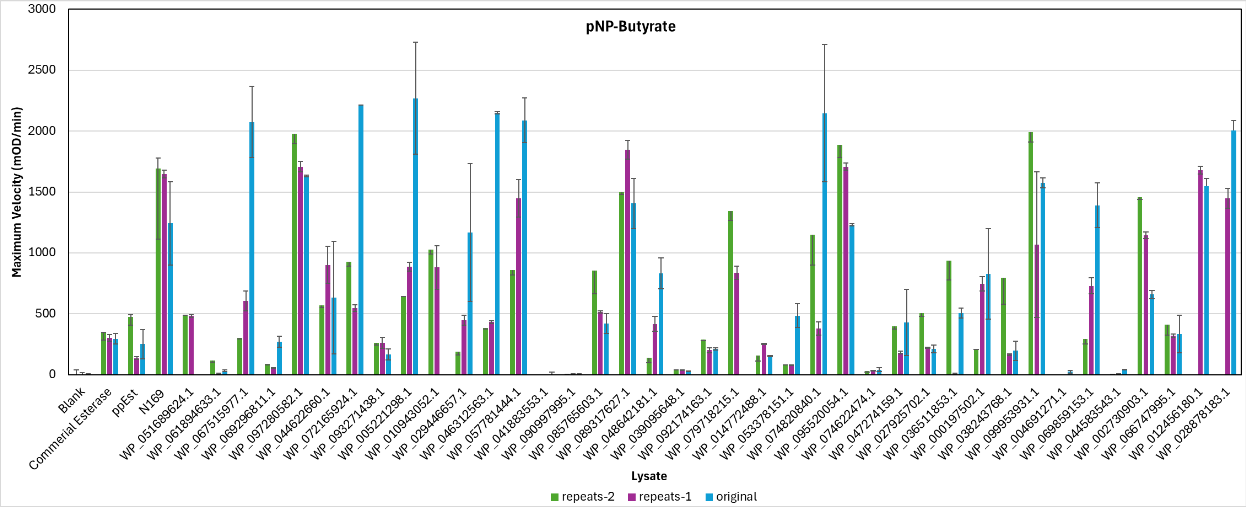
pNP-Butyrate
Maximum Velocity (mOD/min)
3000
2500
2000
1500
1000
500
0
Blank
Commerial Esterase
ppEst
N169
WP_051689624.1
WP_061894633.1
WP_067515977.1
WP_069296811.1
WP_097280582.1
WP_044622660.1
WP_072165924.1
WP_093271438.1
WP_005221298.1
WP_010943052.1
WP_029446657.1
WP_046312563.1
WP_057781444.1
WP_041883553.1
WP_090997995.1
WP_085765603.1
WP_089317627.1
WP_048642181.1
WP_039095648.1
WP_092174163.1
WP_079718215.1
WP_014772488.1
WP_053378151.1
WP_074820840.1
WP_095520054.1
WP_074622474.1
WP_047274159.1
WP_027925702.1
WP_036511853.1
WP_000197502.1
WP_038243768.1
WP_099953931.1
WP_004691271.1
WP_069859153.1
WP_044583543.1
WP_002730903.1
WP_066747995.1
WP_012456180.1
WP_028878183.1
Lysate
repeats-2
repeats-1
original

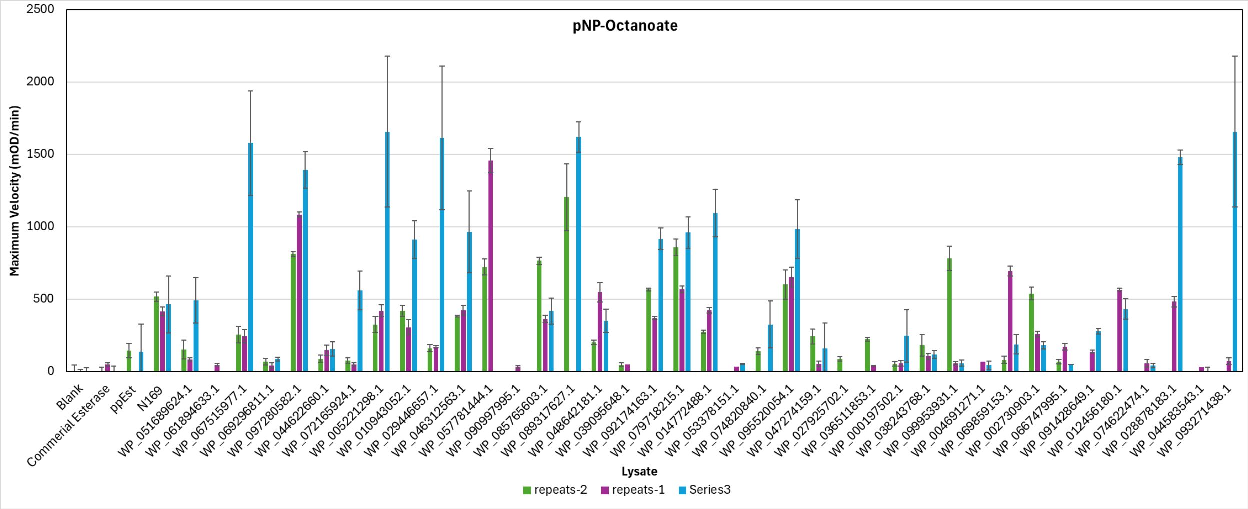
pNP-Octanoate
Maximum Velocity (mOD/min)
2500
2000
1500
1000
500
0
Blank
Commercial Esterase
ppEst
N169
WP_051689624.1
WP_061894633.1
WP_067515977.1
WP_069296811.1
WP_097280582.1
WP_044622660.1
WP_072165924.1
WP_005221298.1
WP_010943052.1
WP_029446657.1
WP_046312563.1
WP_057781444.1
WP_090997995.1
WP_085765603.1
WP_089317627.1
WP_048642181.1
WP_039095648.1
WP_092174163.1
WP_079718215.1
WP_014772488.1
WP_053378151.1
WP_074820840.1
WP_095520054.1
WP_047274159.1
WP_027925702.1
WP_036511853.1
WP_000197502.1
WP_038243768.1
WP_099953931.1
WP_004691271.1
WP_069859153.1
WP_002730903.1
WP_066747995.1
WP_091428649.1
WP_012456180.1
WP_074622474.1
WP_028878183.1
WP_044583543.1
WP_093271438.1
Lysate
repeats-2
repeats-1
Series3

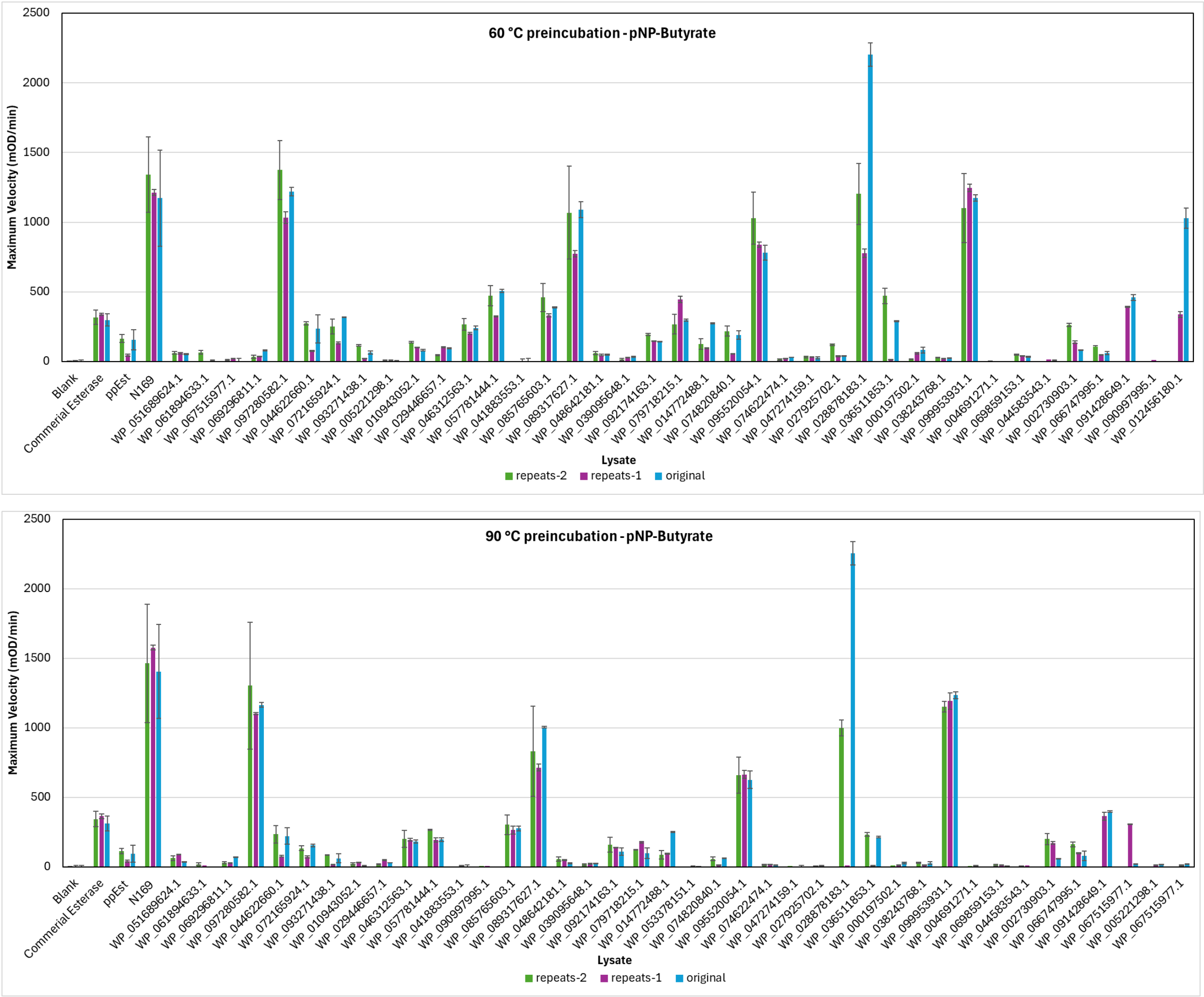


**Figure S4. Reproducibility of enzyme activity measurements across screening plates.** Activity measurements were repeated for lysates of a subset of 40 enzymes to evaluate reproducibility of the high-throughput screening workflow. Plots compare maximum velocity (MaxV, mOD/min) values obtained in the initial screen and in independent repeat measurements across assay conditions. Error bars represent the standard deviation (n = 3 on average).

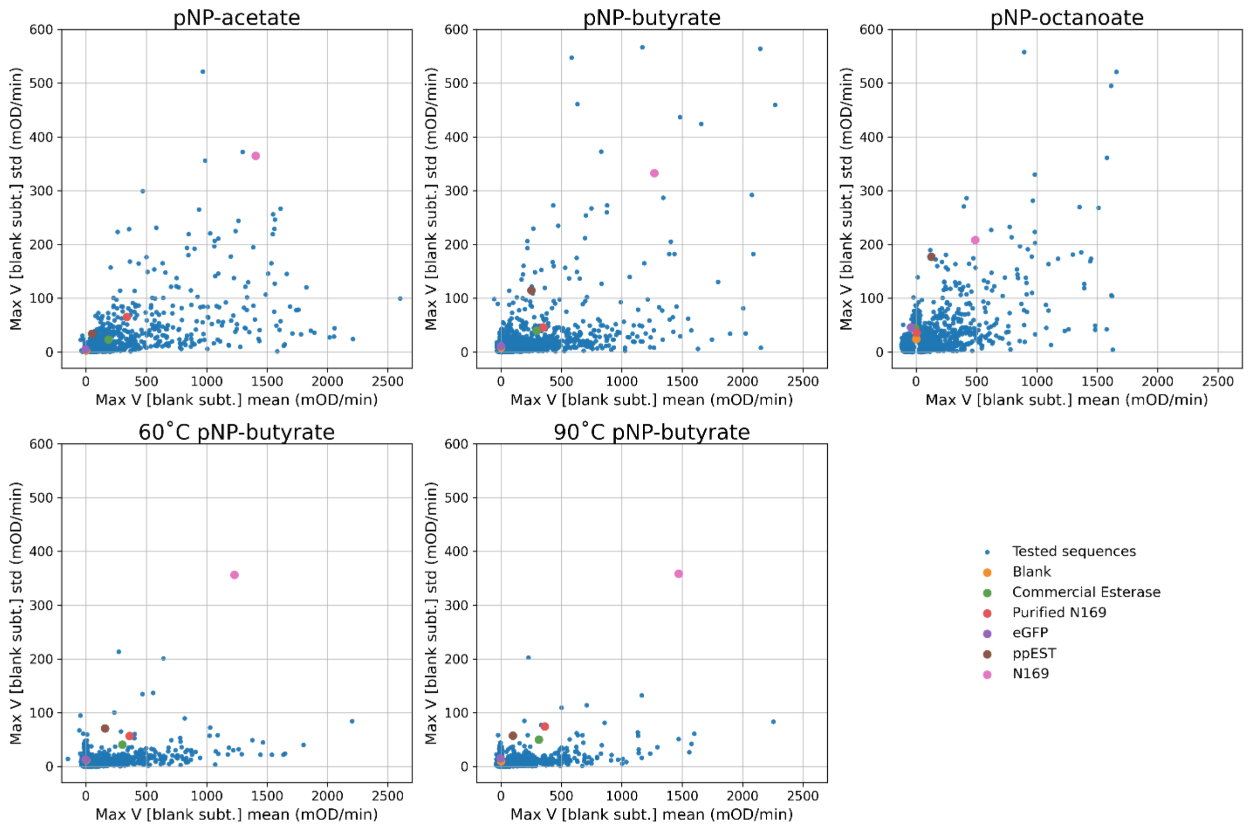


**Figure S5. Mean-variance relationship of high-throughput lysate assays.** Scatterplots comparing the mean (x-axis) against the standard deviation (y-axis) of replicate measurement of maximum velocity for each enzyme variant. Data are shown for the three substrates (pNP-acetate, pNP-butyrate, pNP-octanoate) and the two heat-treatment conditions (pNP-butyrate at 60°C and 90°C). While variance tends to increase with higher activity, the majority of points cluster near the X-axis, indicating consistent reproducibility across the high-throughput lysate screening campaign. Reference controls (blank, eGFP, commercial esterase, lysates for PpEST and N169 (unpublished) and purified N169) are highlighted.

OFFICIAL

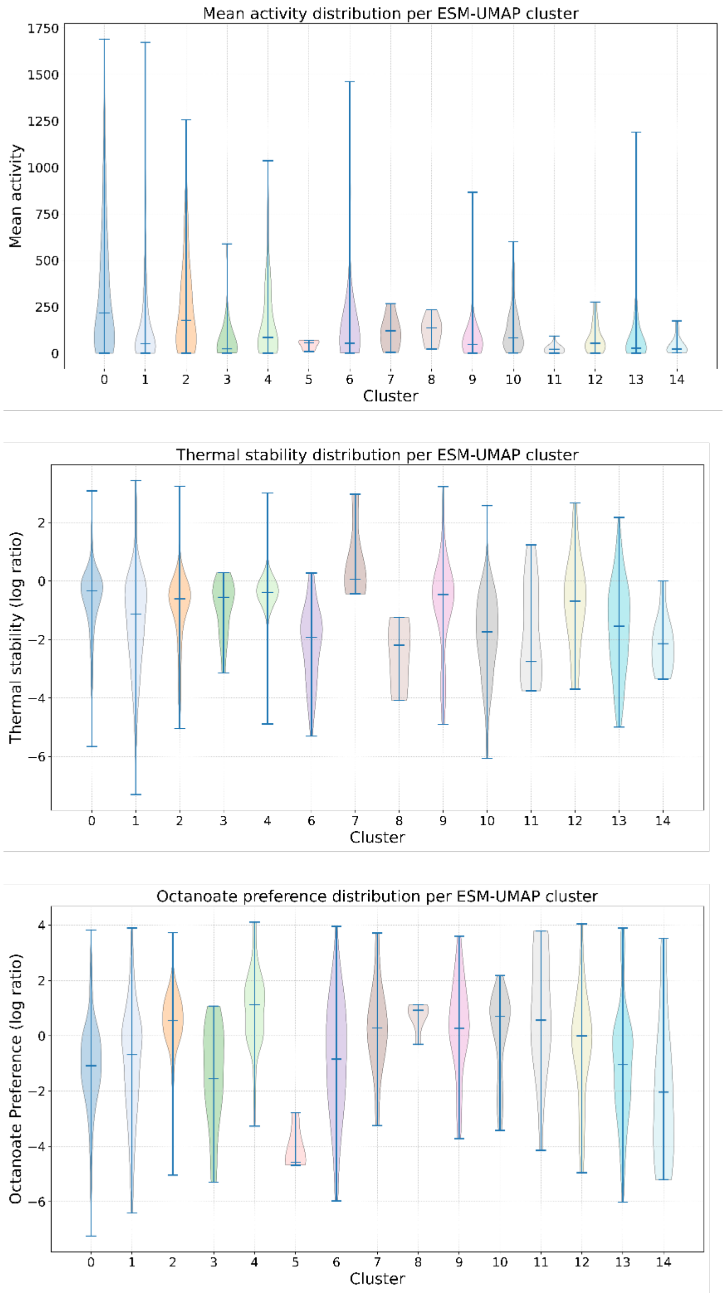


**Figure S6. Distribution of functional properties across sequence clusters.** Violin plots showing the distribution of experimentally measured functional properties for the 15 spectral clusters defined by the ESM-UMAP analysis. The clusters and colours correspond to Figure 2. Horizontal bars within violins indicate the extrema and median values. (Top) Mean activity distributions reveal that while most variants exhibit moderate activity, clusters 0, 1, 2, 4, 6, and 13 contain rare, high-activity outliers. (Middle) Thermal stability (log retention ratio) varies by cluster, with Cluster 7 displaying the highest median stability and Cluster 8 the lowest. (Bottom) Octanoate preference highlights functional divergence; Cluster 5 is strictly acetate-preferring (negative log ratio), while Cluster 4 shows a median preference for octanoate.

OFFICIAL

OFFICIAL

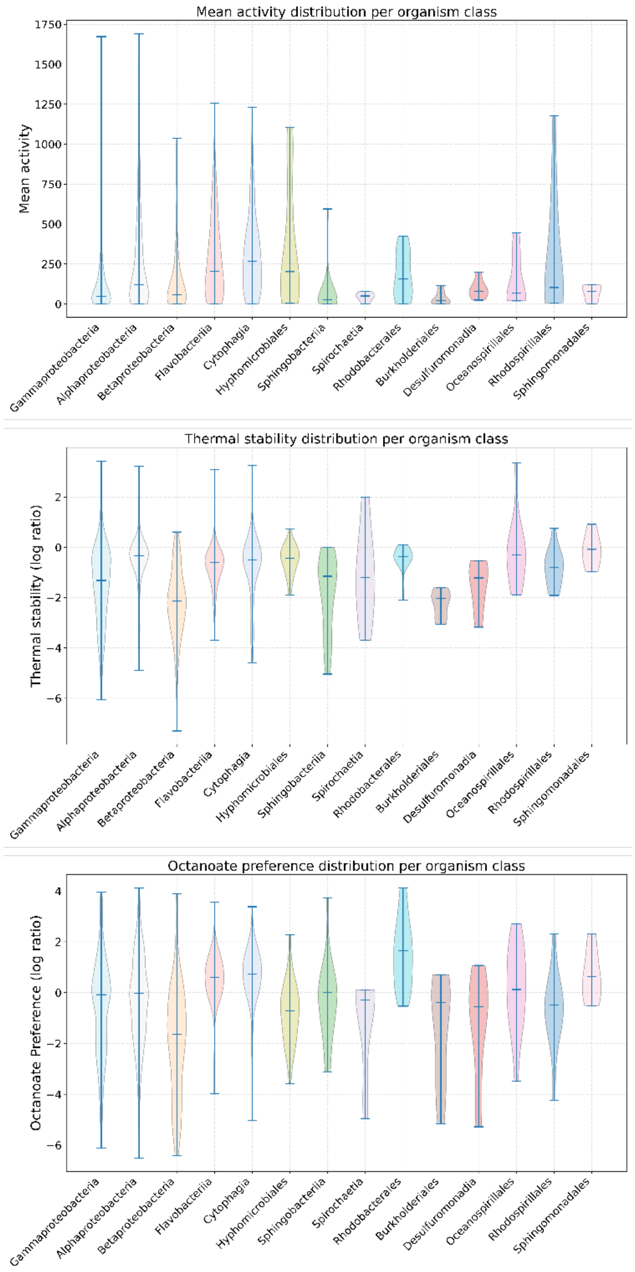


**Figure S7. Distribution of functional properties across taxonomic groups.** Violin plots showing the distribution of experimentally measured functional properties grouped by the taxonomic classification of the source organism. The colours correspond to Figure 2. Horizontal bars within violins indicate the extrema and median values. **(Top)** Mean activity distributions highlight that specific orders such as Hyphomicrobiales and classes like Cytophagia generally exhibit higher activity than the broader Alpha- or Gamma proteobacteria classes. **(Middle)** Thermal stability is taxonomically stratified; Sphingomonadales displays the highest median stability, whereas Betaproteobacteria are notably heat-sensitive. **(Bottom)** Octanoate preference reveals lineage-specific preferences, with Rhodobacterales showing a distinct preference for octanoate (positive log ratio) and Spirochaetia showing a distinct preference for acetate (negative log ratio).

OFFICIAL

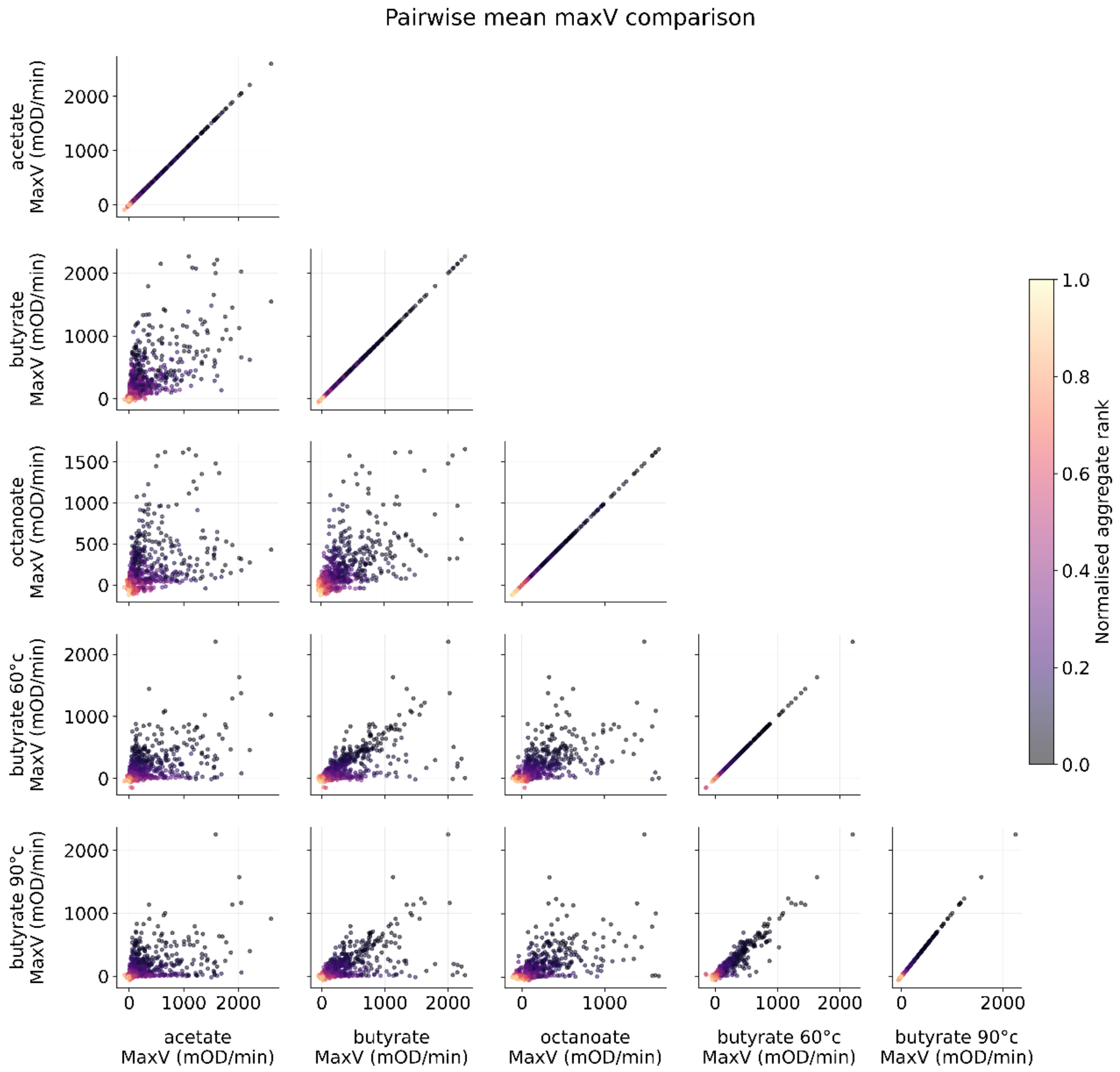


**Figure S8. Pairwise comparisons of experimental activity.** Scatterplot matrix comparing the relationships between maximum velocity (MaxV, mOD/min), of the 1,513 sequences in the initial screen, measured across the three substrates (pNP-acetate, pNP-butyrate, pNP-octanoate) and two thermal treatments (60°C and 90°C). Points are coloured by their normalised aggregate rank (the target metric for overall activity), where darker values indicate higher aggregate performance.

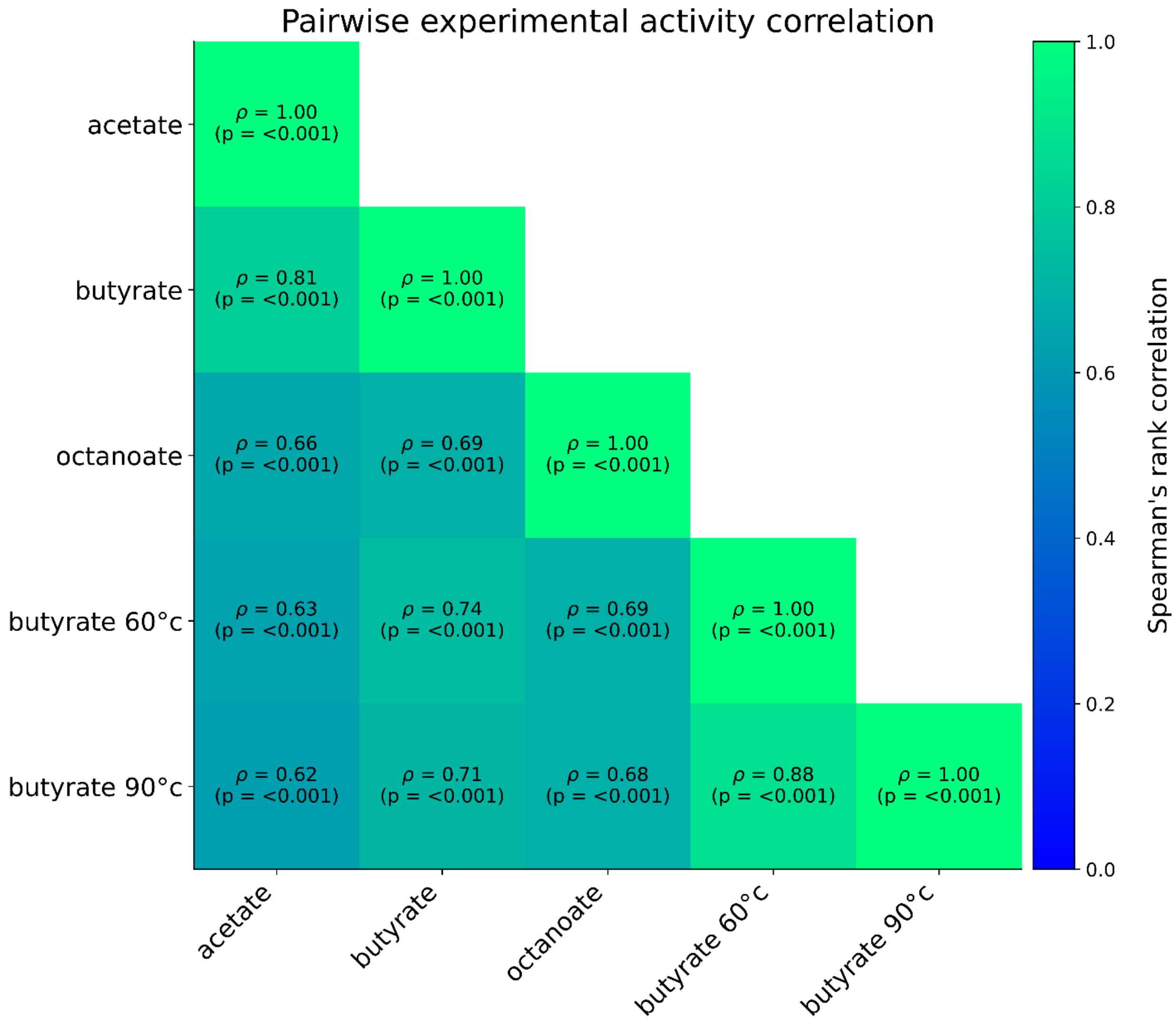


**Figure S9. Pairwise experimental activity correlation.** Heatmap displaying Spearman's rank correlation coefficients (ρ) between experimentally measured activities (MaxV) across three substrates (acetate, butyrate, octanoate) and two thermal treatments (60°C and 90°C) for the initial training cohort. All calculated correlations are highly significant ($p<0.001$). Activity on short-chain substrates (acetate and butyrate) is strongly correlated ($\rho=0.81$), whereas the correlation between the shortest and longest chains (acetate and octanoate) is notably weaker ($\rho=0.66$), reflecting divergent chain-length specificities across the enzyme family. Additionally, the strong correlation between the 60°C and 90°C thermal treatments ($\rho=0.88$) confirms the consistency of the heat-shock assay.

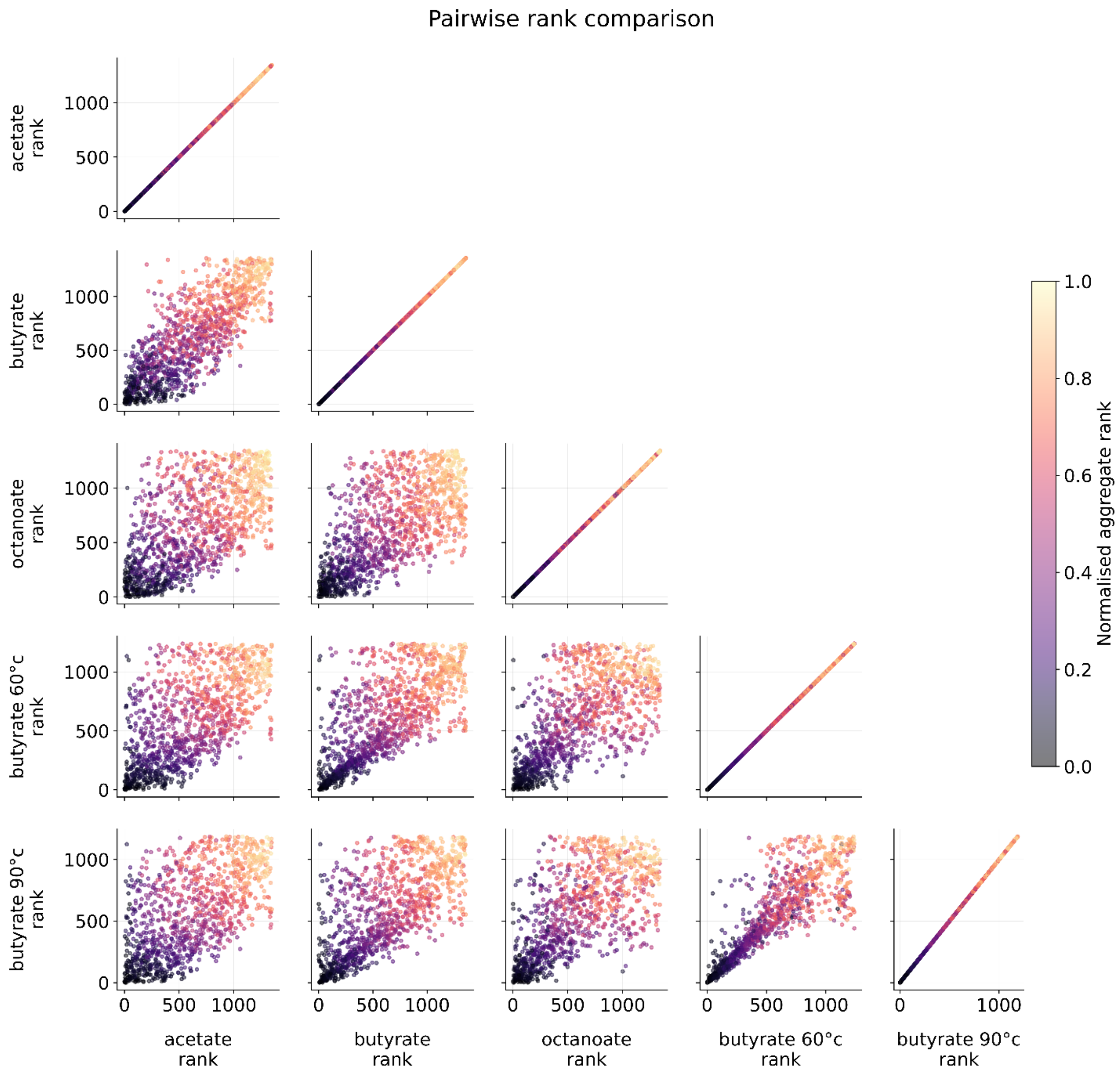


**Figure S10. Pairwise comparisons of experimental activity ranks (lower is better).** Scatterplot matrix comparing the relative ranking of the 1,513 sequences in the initial screen across the three substrates (pNP-acetate, pNP-butyrate, pNP-octanoate) and two thermal treatments (60°C and 90°C). Points are coloured by their normalised aggregate rank (the target metric for overall activity), where darker values indicate higher aggregate performance. The tight correlation between acetate and butyrate ranks contrasts with the broader dispersion observed for octanoate, reinforcing that long-chain specificity is a distinct functional trait from short-chain activity.

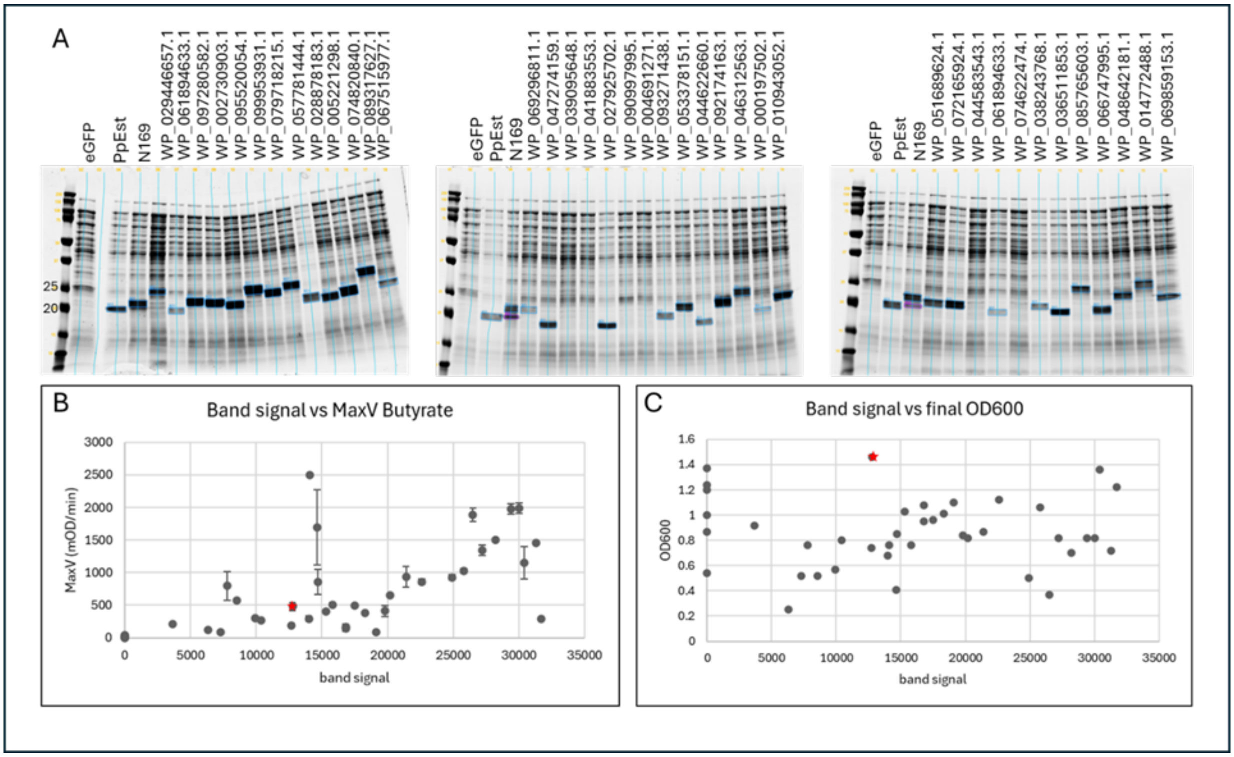


**Figure S11.** Lysate from thirty-nine repeats with a range in activity (Figure S4, repeats-2), were run on SDS-page to examine protein expression. 18 µl of a 1/10 dilution of the lysate for each sample was run in a total of 30 µl (1x Laemmli sample buffer and Bio-Rad XT reducing agent) on one of three 12% tris-glycine (Bio-Rad Criterion™ TGX™ Precast Midi Protein Gel, 18 well) gels alongside eGFP, ppEST and N169. The gels were run at 200V for 50 min in a Bio-Rad Criterion gel with 1x tris-glycine buffer and stained with Bulldog Aquastain (Geneworks). The gels were scanned on the Li-Cor Odyssey on channel 700 with the same settings and analysed with Empiria Studio® Software (version 3.3.0.195) as shown in A. Esterase bands were identified by size and comparison to controls (highlighted by blue boxes on gel photos) and Empiria used to measure the band signal which was compared to the average MaxV for pNP-Butyrate (B) and final culture OD600 (C). PpEst is indicated by a red star. In most cases expression appeared to approximately correlate with activity but there were some exceptions. In all cases, when no esterase band was visible on the gel, the activity was very low or zero. There did not appear to be a relationship between culture final OD600 and expression.

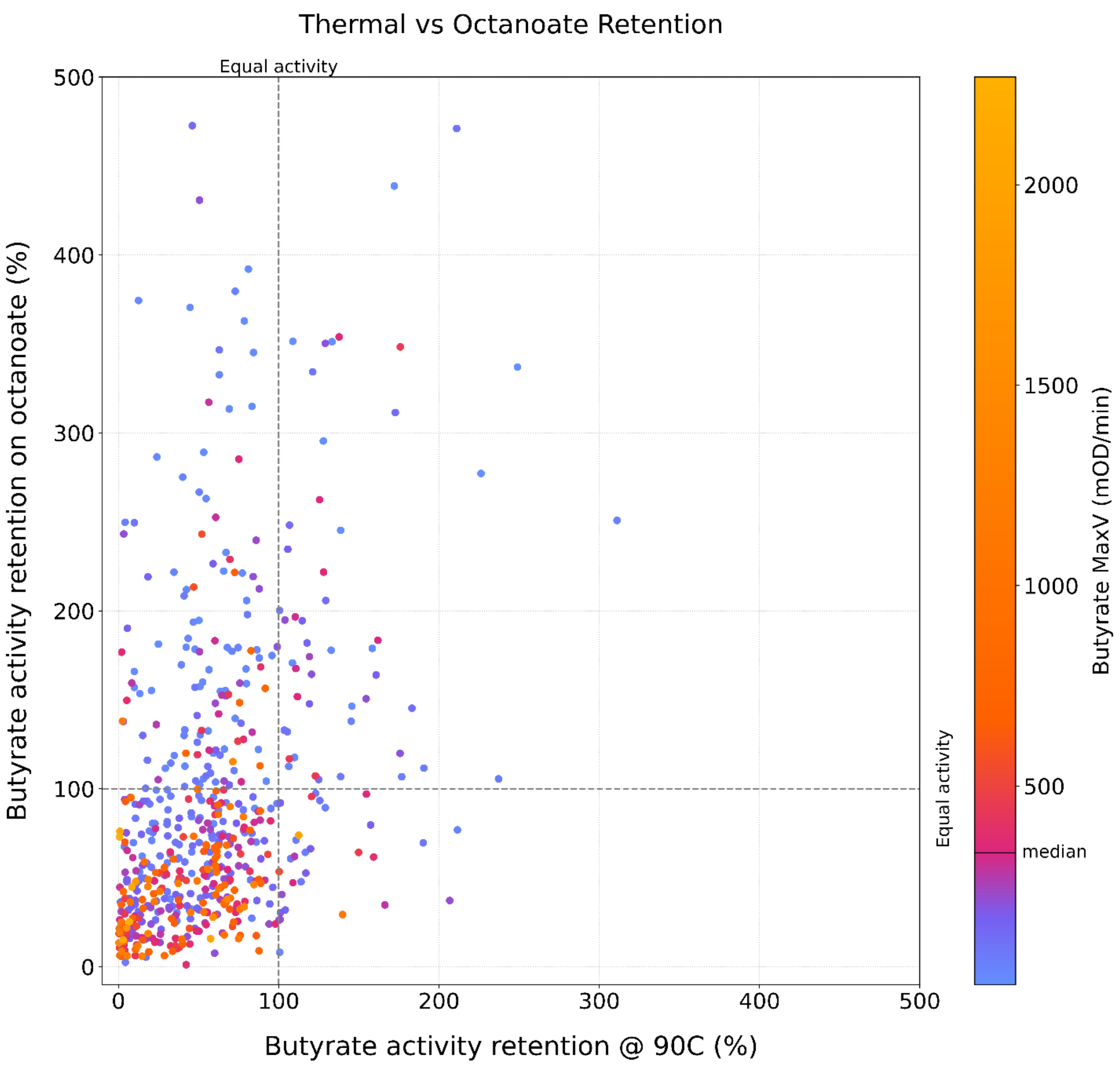


**Figure S12. Relationship between thermal stability and substrate preference.** Scatterplot displaying the correlation between thermal resilience (X-axis, retained butyrate activity after 90°C treatment) and octanoate preference (Y-axis, octanoate activity as a percentage of butyrate activity) for the 1,513 sequences in the initial screen. Points are coloured by butyrate activity (MaxV), with the median activity level marked on the colour bar. Dashed lines indicate activity equivalence. The scatted relationship between the points at high retention values suggests that thermal stability and a preference for octanoate are largely independent traits. Notably, the variants with highest pNP-butyrate activity (bright orange) cluster in the low thermal stability and low octanoate preference region. Variants with high thermal stability or octanoate preference tend to exhibit lower absolute catalytic rates on butyrate, suggesting a potential trade-off between specialisation and peak catalytic turnover.

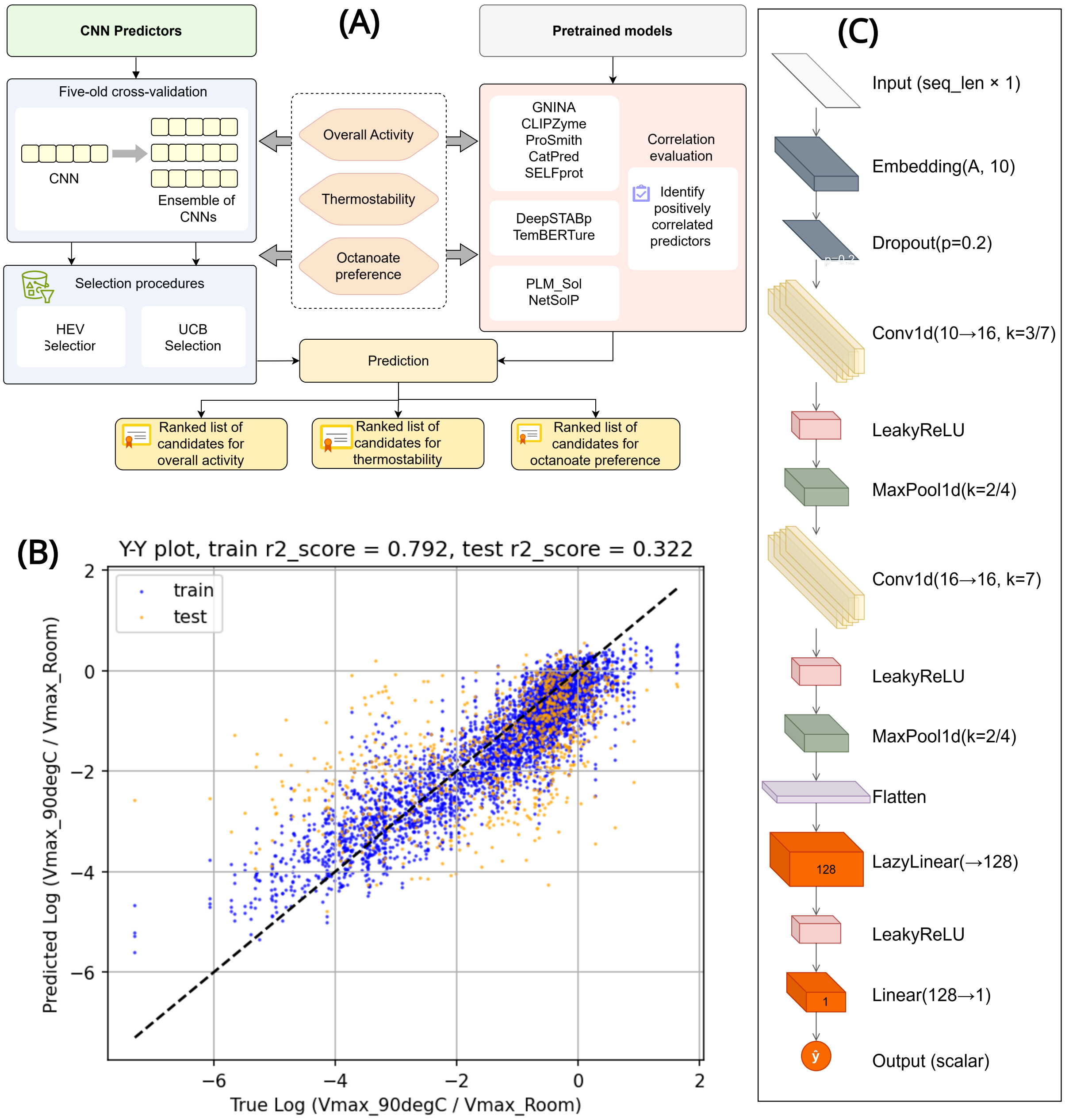


**Figure S13.** Model development and validation. **A.** Schematic representation of workflow for model building, training, validation and prediction. **B.** CNN architecture. **C.** Y-Y plot of the CNN ensemble's aggregate predictions vs. the true thermostability target, $y_{TS}$, for the in-fold (blue) and out-of-fold (orange) predictions.

**Table S2. Spearman's correlations between predictions from pre-trained models and experimentally measured data.**

| | Dataset | | | | | |
|---|---|---|---|---|---|---|
| **Pre-trained model** | **All substrates** | **Acetate only** | **Butyrate Only** | **Octanoate Only** | **Substrate Specificity** | **Thermostability** |
| CLIPZyme[2] | -0.122 | 0.072 | -0.006 | -0.105 | -0.275 | |
| ESP ProSmith[3] | -0.034 | -0.010 | -0.021 | 0.013 | -0.040 | |
| CatPred[4] $\log_{10}(k_{cat})$ max | 0.193 | 0.103 | 0.148 | 0.216 | 0.217 | |
| CatPred[4] $\log_{10}(K_m)$ mean* | 0.128 | 0.150 | 0.081 | 0.177 | 0.087 | |
| SELFProt[5] $\log_{10}(k_{cat})$ | -0.111 | -0.085 | 0.003 | -0.166 | -0.212 | |
| SELFProt[5] $\log_{10}(K_m)$* | 0.051 | 0.022 | 0.001 | 0.019 | 0.065 | |
| Netsolp[6] solubility | 0.214 | 0.141 | 0.184 | 0.332 | 0.368 | |
| PLM Sol[7] | 0.075 | 0.095 | 0.128 | 0.012 | 0.011 | |
| GNINA[8] Vina affinity score* | 0.107 | 0.051 | 0.098 | 0.090 | 0.073 | |
| GNINA[8] CNN VS score | -0.051 | -0.181 | -0.012 | 0.111 | 0.163 | |
| GNINA[8] CNN affinity score | -0.075 | -0.101 | 0.052 | 0.015 | -0.015 | |
| DeepStabP[9] $T_m$ | | | | | | 0.139 |
| TemBERTure[10] $T_m$ | | | | | | 0.071 |
| Aggregate rank all substrates (catpredkcat, netsolp) | 0.283 | | | | | |
| Aggregate rank octanoate only (catpredkcat, netsolp, GNINA CNN VS)* | | | | | -0.410 | |

*More negative Spearman's correlation coefficients indicate stronger agreement with experimental measurements.

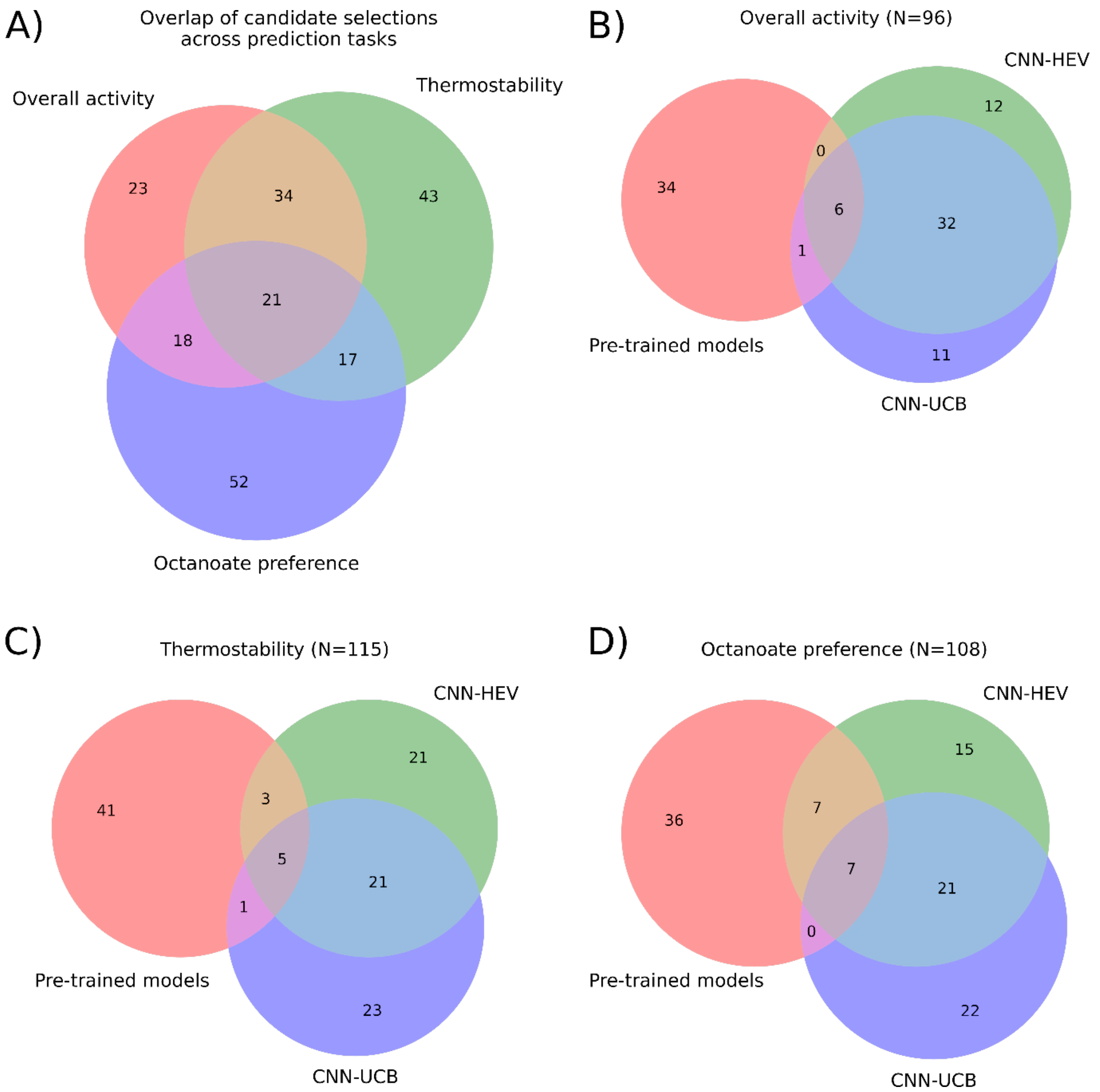


**Figure S14. Composition and overlap of the model-guided validation panel.** Venn diagrams showing the breakdown of the 200 model-selected sequences within the 230-sequence validation cohort (random selections excluded). **(A)** Overlap of candidate selections across the three prediction tasks. The central intersection (n=21) represents sequences prioritised for all three functional objectives, indicating convergence on shared sequence features across tasks. **(B-D)** Overlap of sequences prioritised by pre-trained models, CNN-HEV, and CNN-UCB strategies for **(B)** overall activity, **(C)** thermostability, and **(D)** octanoate preference. High agreement is observed between the two CNN strategies, which share a significant core of candidates, whereas pre-trained models prioritise a largely distinct set of candidates.

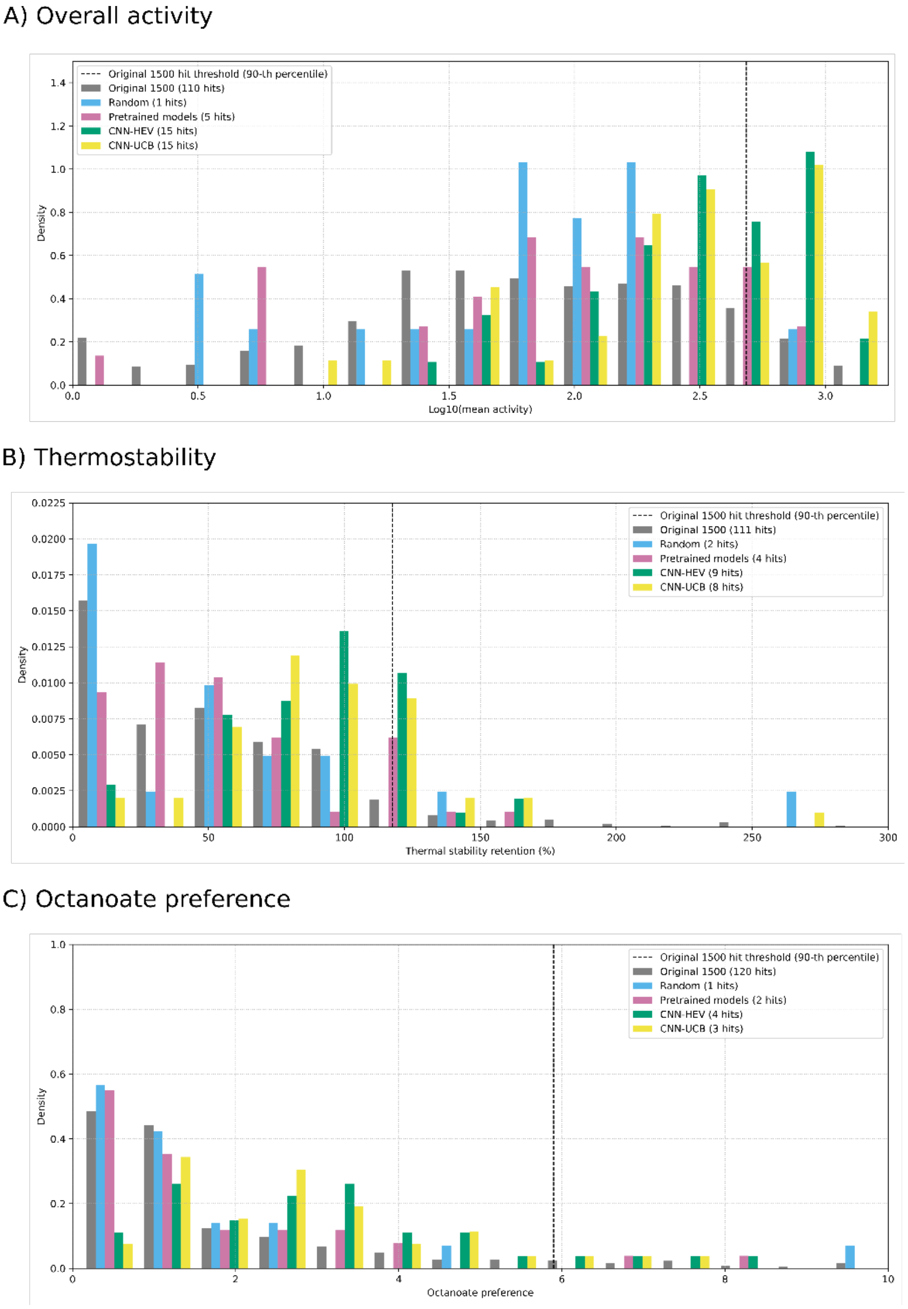


**Figure S15. Experimental validation of ML-guided sequence selection.** Histograms comparing the experimentally measured performance of the original screen (grey) against prospective validation panels selected by random sampling (blue), pretrained models (pink), CNN-HEV (green) and CNN-UCB (yellow) strategies. The vertical dashed line marks the 90th percentile "hit" threshold established by the original screen. The total number of "hits" for each policy is listed in the legend. **(A)** Overall activity results show an enrichment of high-activity variants by both CNN strategies, with distributions heavily skewed toward the upper quantiles compared to random and pretrained baselines. **(B)** Thermostability retention (%) distributions demonstrate that CNN-guided selections shift the population toward higher stability. **(C)** Octanoate preference distributions show that CNN strategies enriched for sequences with a high preference for octanoate substrates, retrieving a greater number of hits above the 90th-percentile threshold compared to the random or pretrained baselines.

**Table S3. Statistical evaluation of functional improvement across machine learning selection strategies.** To evaluate whether a selection strategy successfully shifted the overall quality of the baseline experimental library (original screen), a one-sided Mann-Whitney U test was performed. The difference in medians quantifies the magnitude of this shift. The data are partitioned by the three tasks: (A) Mean activity, (B) Thermostability, and (C) Octanoate preference. For all tasks, both CNN strategies demonstrated highly statistically significant positive distributional shifts ($p < 0.001$) compared to the original screening baseline, whereas the pre-trained models and random selections failed to provide significant improvement.

A) Mean activity

| Data | Median | Median shift | p (Mann-Whitney) |
|---|---|---|---|
| Original screen | 1.831744 | - | - |
| Second screen | 2.188494 | 0.356749 | 0.000001 |
| Random | 1.832969 | 0.001225 | 0.629449 |
| Pre-trained models | 2.090404 | 0.25866 | 0.084164 |
| CNN-HEV | 2.560192 | 0.728447 | 0.0 |
| CNN-UCB | 2.461449 | 0.629704 | 0.0 |

B) Thermostability

| Data | Median | Median shift | p (Mann-Whitney) |
|---|---|---|---|
| Original screen | 47.740815 | - | - |
| Second screen | 61.039939 | 13.299124 | 0.000612 |
| Random | 47.175076 | -0.565739 | 0.441468 |
| Pre-trained models | 49.302254 | 1.561439 | 0.256418 |
| CNN-HEV | 88.090940 | 40.350125 | 0.0 |
| CNN-UCB | 86.771366 | 39.030551 | 0.0 |

C) Octanoate preference

| Data | Median | Median shift | p (Mann-Whitney) |
|---|---|---|---|
| Original screen | 1.000000 | - | - |
| Second screen | 0.659802 | -0.340198 | 0.920628 |
| Random | 1.000000 | 0.0 | 0.631904 |
| Pre-trained models | 0.849793 | -0.150207 | 0.666177 |
| CNN-HEV | 2.729245 | 1.729245 | 0.000012 |
| CNN-UCB | 2.389113 | 1.389113 | 0.00001 |

**Table S4. Statistical evaluation of discovery enrichment across machine learning selection strategies.** To evaluate whether a selection strategy successfully enriched for top-tier functional variants compared to the baseline experimental library (original screen), a one-sided Fisher's Exact Test was performed. A "hit" was defined as any sequence exceeding the 90th percentile performance of the original screen. The Enrichment Factor (EF) quantifies the ratio of the new hit rate relative to the baseline. The data are partitioned by the three tasks: (A) Mean activity, (B) Thermostability, and (C) Octanoate preference. For overall activity (A), both supervised CNN strategies demonstrated highly statistically significant, multi-fold enrichment ($p < 0.001$). For thermostability and octanoate preference (B and C), no selection strategies achieved statistically significant enrichment at this strict threshold, highlighting the difficulty of predicting extreme outliers for these specific traits.

A) Mean activity

| Data | N | Hits | Hit rate | EF [CI95 low, CI95 high] | p |
|---|---|---|---|---|---|
| Original screen | 1094 | 110 | 0.100548 | - | - |
| Second screen | 178 | 30 | 0.168539 | 1.6762 [1.156282, 2.429898] | 0.007119 |
| Random | 18 | 1 | 0.055556 | 0.552525 [0.081575, 3.742352] | 0.851671 |
| Pre-trained models | 34 | 5 | 0.147059 | 1.462567 [0.63859, 3.349724] | 0.260538 |
| CNN-HEV | 43 | 15 | 0.348837 | 3.469345 [2.222873, 5.414772] | 0.000019 |
| CNN-UCB | 41 | 15 | 0.365854 | 3.638581 [2.342809, 5.651025] | 0.00001 |

B) Thermostability

| Data | N | Hits | Hit rate | EF [CI95 low, CI95 high] | p |
|---|---|---|---|---|---|
| Original screen | 1110 | 111 | 0.100000 | - | - |
| Second screen | 186 | 22 | 0.118280 | 1.182796 [0.769235, 1.818696] | 0.259633 |
| Random | 19 | 2 | 0.105263 | 1.052632 [0.28042, 3.951335] | 0.582281 |
| Pre-trained models | 45 | 4 | 0.088889 | 0.888889 [0.343111, 2.302819] | 0.672497 |
| CNN-HEV | 48 | 9 | 0.187500 | 1.875 [1.013932, 3.467318] | 0.052035 |
| CNN-UCB | 47 | 8 | 0.170213 | 1.702128 [0.883771, 3.278267] | 0.100541 |

C) Octanoate preference

| Data | N | Hits | Hit rate | EF [CI95 low, CI95 high] | p |
|---|---|---|---|---|---|
| Original screen | 1199 | 120 | 0.100083 | - | - |
| Second screen | 190 | 5 | 0.026316 | 0.262939 [0.108909, 0.634812] | 0.999955 |
| Random | 23 | 1 | 0.043478 | 0.43442 [0.063411, 2.976177] | 0.911191 |
| Pre-trained models | 43 | 2 | 0.046512 | 0.464729 [0.118814, 1.817734] | 0.936573 |
| CNN-HEV | 41 | 4 | 0.097561 | 0.974797 [0.378392, 2.511226] | 0.600276 |
| CNN-UCB | 39 | 3 | 0.076923 | 0.76859 [0.255749, 2.309806] | 0.7627 |

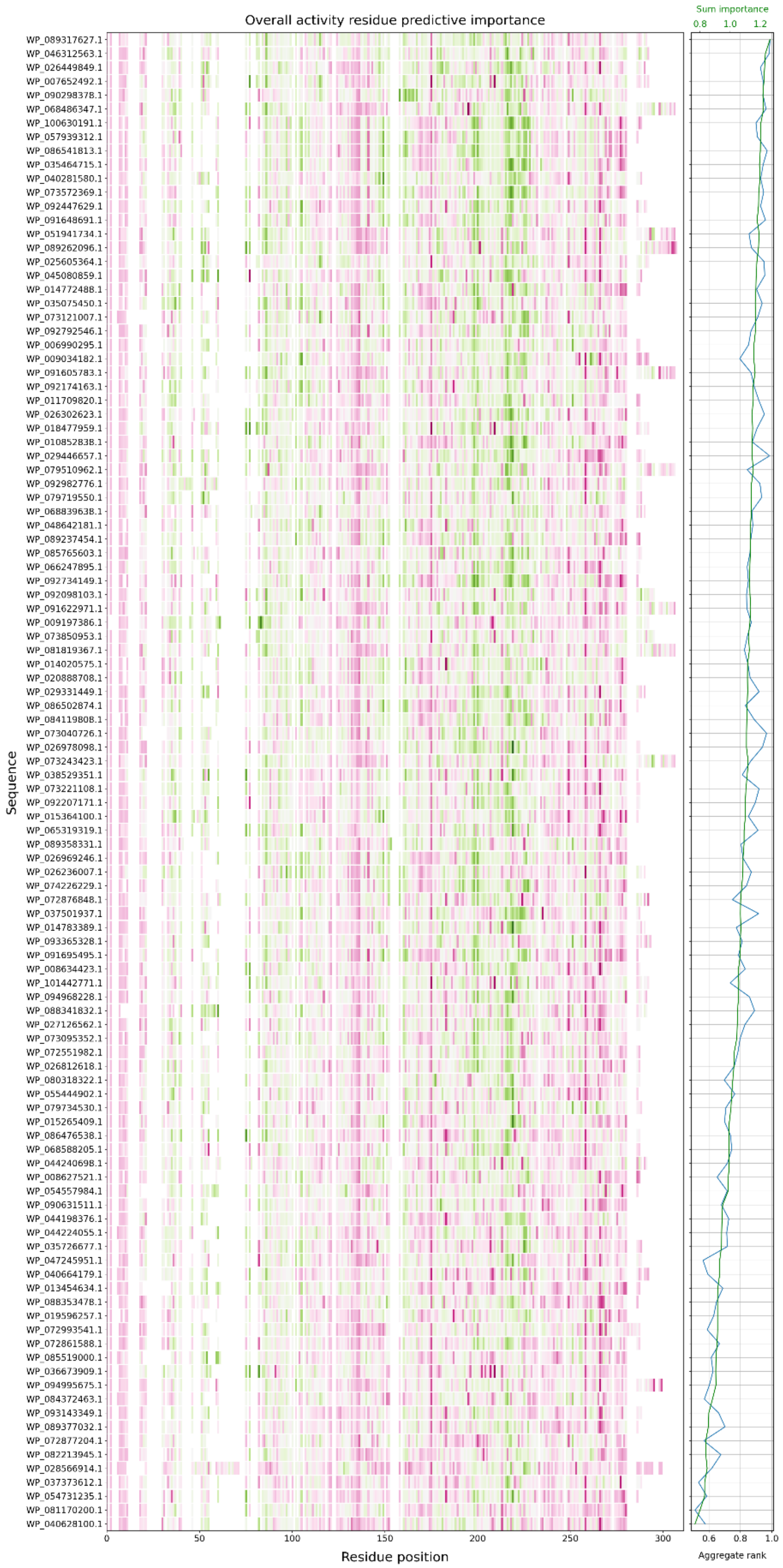
Overall activity residue predictive importance
Sequence
Residue position
0
50
100
150
200
250
300
Sum importance
0.8
1.0
1.2
0.6
0.8
1.0
Aggregate rank

OFFICIAL

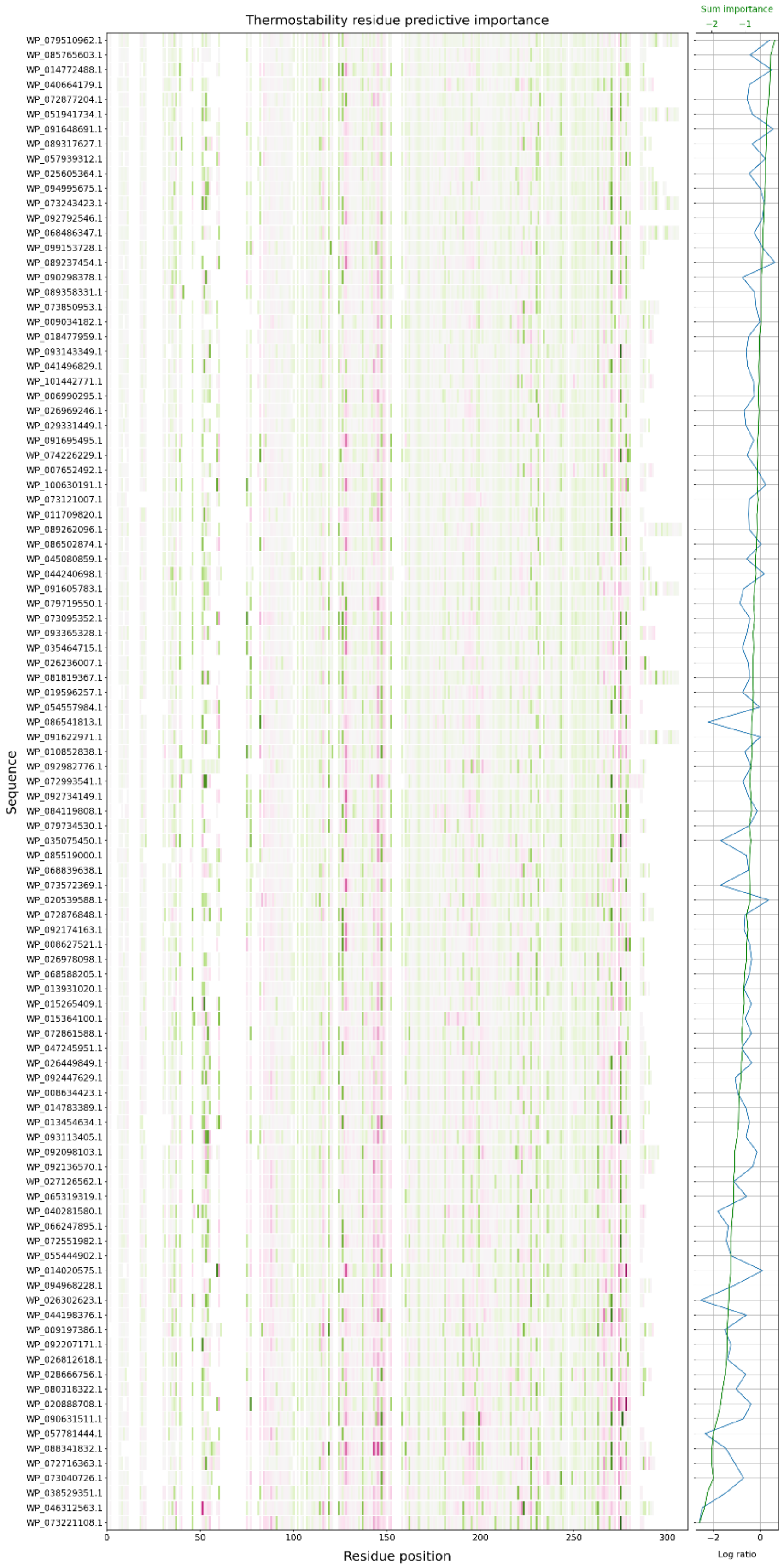


OFFICIAL

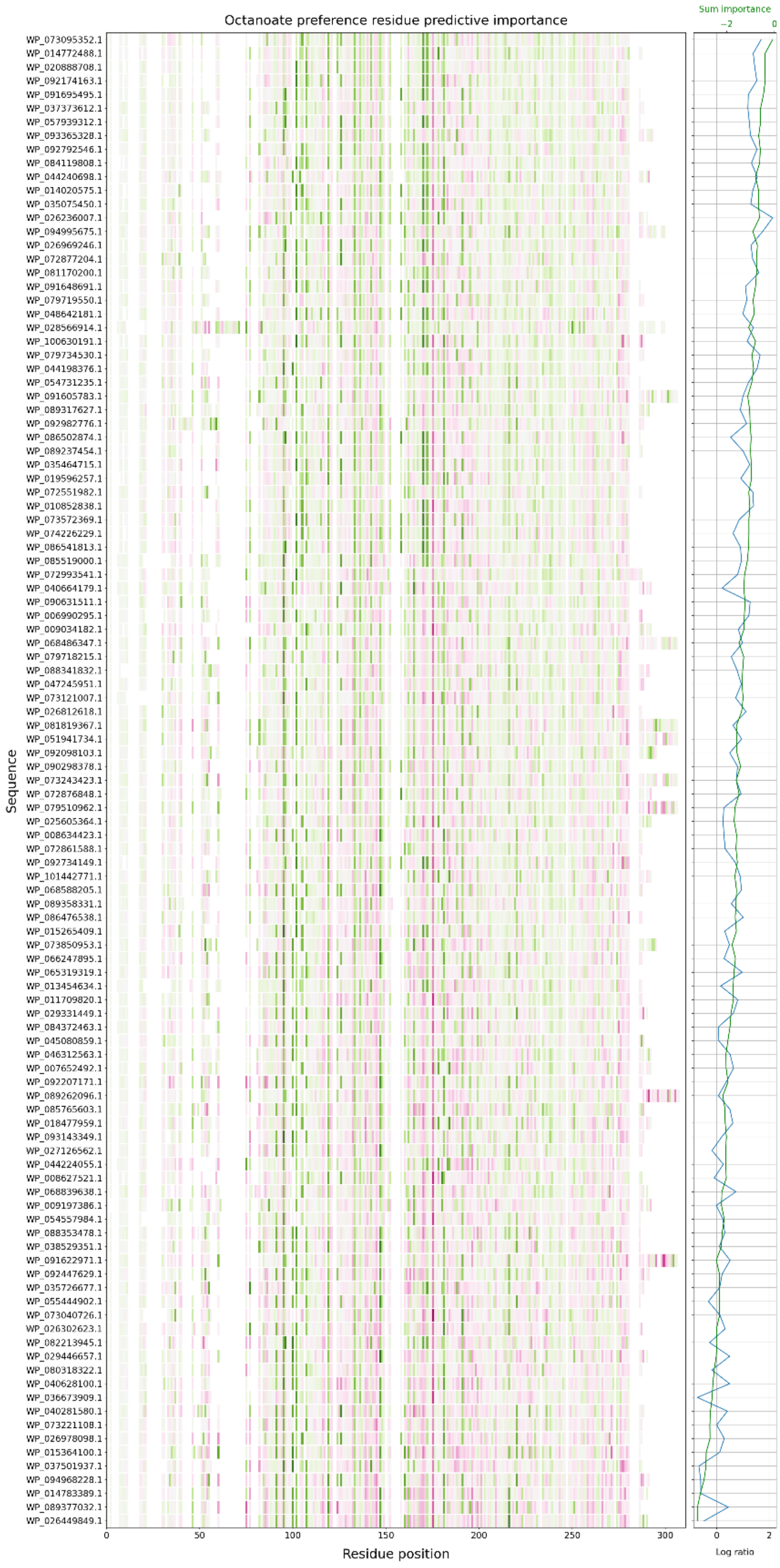
Octanoate preference residue predictive importance
Sum importance
−2
0
Sequence
WP_073095352.1
WP_014772488.1
WP_020888708.1
WP_092174163.1
WP_091695495.1
WP_037373612.1
WP_057939312.1
WP_093365328.1
WP_092792546.1
WP_084119808.1
WP_044240698.1
WP_014020575.1
WP_035075450.1
WP_026236007.1
WP_094995675.1
WP_026969246.1
WP_072877204.1
WP_081170200.1
WP_091648691.1
WP_079719550.1
WP_048642181.1
WP_028566914.1
WP_100630191.1
WP_079734530.1
WP_044198376.1
WP_054731235.1
WP_091605783.1
WP_089317627.1
WP_092982776.1
WP_086502874.1
WP_089237454.1
WP_035464715.1
WP_019596257.1
WP_072551982.1
WP_010852838.1
WP_073572369.1
WP_074226229.1
WP_086541813.1
WP_085519000.1
WP_072993541.1
WP_040664179.1
WP_090631511.1
WP_006990295.1
WP_009034182.1
WP_068486347.1
WP_079718215.1
WP_088341832.1
WP_047245951.1
WP_073121007.1
WP_026812618.1
WP_081819367.1
WP_051941734.1
WP_092098103.1
WP_090298378.1
WP_073243423.1
WP_072876848.1
WP_079510962.1
WP_025605364.1
WP_008634423.1
WP_072861588.1
WP_092734149.1
WP_101442771.1
WP_068588205.1
WP_089358331.1
WP_086476538.1
WP_015265409.1
WP_073850953.1
WP_066247895.1
WP_065319319.1
WP_013454634.1
WP_011709820.1
WP_029331449.1
WP_084372463.1
WP_045080859.1
WP_046312563.1
WP_007652492.1
WP_092207171.1
WP_089262096.1
WP_085765603.1
WP_018477959.1
WP_093143349.1
WP_027126562.1
WP_044224055.1
WP_008627521.1
WP_068839638.1
WP_009197386.1
WP_054557984.1
WP_088353478.1
WP_038529351.1
WP_091622971.1
WP_092447629.1
WP_035726677.1
WP_055444902.1
WP_073040726.1
WP_026302623.1
WP_082213945.1
WP_029446657.1
WP_080318322.1
WP_040628100.1
WP_036673909.1
WP_040281580.1
WP_073221108.1
WP_026978098.1
WP_015364100.1
WP_037501937.1
WP_094968228.1
WP_014783389.1
WP_089377032.1
WP_026449849.1
0
50
100
150
200
250
300
Residue position
0
2
Log ratio

**Figure S16. Residue-level predictive importance across cluster 2.** Heatmaps show per-residue importance values derived from CNN ensemble models for each task (overall activity, thermostability, octanoate specificity) across sequences from cluster 2. Values are un-normalised and sum approximately to the model prediction for each sequence. Positive (green) and negative (pink) values indicate residues that contribute positively or negatively to the predicted output, respectively. The colour scale is centred on the global median importance value across the heat map to emphasise relative differences across sequences and are shown in **Figure 4**. Consistent vertical patterns reflect sequence regions that reproducibly influence model predictions either positively or negatively across the cluster. The right-hand panel shows the summed importance per sequence and how that compares to the predicted MaxV values.

### References


1. Fu, L., Niu, B., Zhu, Z., Wu, S. & Li, W. CD-HIT: accelerated for clustering the next-generation sequencing data. *Bioinformatics* **28**, 3150–3152 (2012).

2. Mikhael, P. G., Chinn, I. & Barzilay, R. CLIPZyme: Reaction-Conditioned Virtual Screening of Enzymes. *Proc. Mach. Learn. Res.* **235**, 35647–35663 (2024).

3. Kroll, A., Ranjan, S. & Lercher, M. J. A multimodal Transformer Network for protein-small molecule interactions enhances predictions of kinase inhibition and enzyme-substrate relationships. *PLoS Comput. Biol.* **20**, e1012100 (2024).

4. Boorla, V. S. & Maranas, C. D. CatPred: a comprehensive framework for deep learning in vitro enzyme kinetic parameters. *Nature Communications 2025 16:1* **16**, 2072- (2025).

5. Wilson, M., Coudrat, T. & Warden, A. SELFprot: Effective and Efficient Multitask Finetuning Methods for Protein Parameter Prediction. *J. Chem. Inf. Model.* **65**, 3226–3238 (2025).

6. Thumuluri, V. *et al.* NetSolP: predicting protein solubility in Escherichia coli using language models. *Bioinformatics* **38**, 941–946 (2022).

7. Zhang, X. *et al.* PLM_Sol: predicting protein solubility by benchmarking multiple protein language models with the updated Escherichia coli protein solubility dataset. *Brief. Bioinform.* **25**, (2024).

8. McNutt, A. T. *et al.* GNINA 1.0: molecular docking with deep learning. *Journal of Cheminformatics 2021 13:1* **13**, 43- (2021).

9. Jung, F., Frey, K., Zimmer, D. & Mühlhaus, T. DeepSTABp: A Deep Learning Approach for the Prediction of Thermal Protein Stability. *International Journal of Molecular Sciences 2023, Vol. 24,* **24**, (2023).

10. Rodella, C., Lazaridi, S. & Lemmin, T. TemBERTure: advancing protein thermostability prediction with deep learning and attention mechanisms. *Bioinformatics Advances* **4**, (2024).
